# Classical baselines outperform released deep-learning ITS classifiers, which collapse on ITS2 where predictions follow the flanking regions

**DOI:** 10.64898/2026.07.29.741510

**Authors:** Aaron O’Brien, Auguste Gardette, César Marín, Pilar Parada

## Abstract

**Background:** Deep-learning classifiers for the fungal internal transcribed spacer (ITS) report accuracies above 90% and are increasingly proposed for environmental metabarcoding. That application differs from the benchmarks in two ways: surveys sequence a primer-bounded subregion, most often ITS2, rather than the full-length reference sequences the models were trained and tested on, and much of what they recover belongs to genera absent from any reference. We tested whether the reported accuracies transfer.

**Design:** Five methods were scored on the same 5,222 queries, one sequence per genus, at two loci: the full-length ITS record, and the ITS2 subregion of that identical record. The methods are the two pretrained deep-learning classifiers distributed with MycoAI, a convolutional network and a transformer sharing a training corpus of 5.23M sequences; two reference-based methods, best-hit alignment and SINTAX; and HiTaC, a hierarchical logistic regression over *k*-mer counts, which we fitted ourselves to the reference the other two consult. Queries were stratified by whether the query’s genus lies in the pretrained models’ label space, recovered from the distributed models themselves. A *novel* genus is one outside that label space, and outside the reference by the same rule, so no method here can return its correct name; *seen* genera are the rest (§2.2).

**Results:** The design favours the pretrained models, whose queries come from the public dataset they were trained on. Even so, on full-length ITS the other three methods exceed both of them at every rank and in both strata, best-hit alignment recovering the correct family for 92.3% of seen-genus queries against 77.9% and 76.5%. Restricting the identical records to ITS2 costs the reference-based methods under four percentage points of seen-genus family accuracy and costs the two pretrained models 49.6 and 58.0, reducing them to 28.3% and 18.5%; HiTaC refitted at that locus recovers 89.2%, so neither learned classification nor the amplicon is what fails. An ablation identifies the cause. Grafting each query’s unaltered ITS2 between the flanking regions of a donor record from a different phylum returns the *donor’s* family for 34.4% of queries in the convolutional model and 63.7% in the transformer, against the query’s own for 4.2% and 0.8%, from a baseline near zero where no donor is present. The predictions therefore follow the flanking regions rather than the ITS2 barcode, which accounts for the collapse and predicts the same failure for any subregion amplicon. Compounding this, on ITS2 the classifiers’ confidence score all but ceases to separate novel from seen genera (AUROC 0.541 and 0.503, the latter at chance, against 0.866 for alignment identity), and the convolutional model is in addition substantially overconfident there, so for that model the failure is not detectable from its own output at all.

**Recommendation:** Reported accuracies for such models should specify the amplicon region of the evaluation, state the length distribution of the training corpus, and be accompanied by a same-query classical baseline.

## 1 Introduction

Metabarcoding of the fungal ITS region routinely recovers sequences with no close match in curated references such as UNITE [17], environmental “dark matter” novel at the genus level or below, and in many surveys such sequences are the majority of what is recovered [23]. Two features of that material set the problem this paper addresses. It is taxonomically unfamiliar, in the strong sense that the correct name may be absent from every reference a method could consult. And it arrives as a primer-bounded subregion, most often ITS2, because that is what survey protocols amplify, rather than as the full-length ITS record that reference databases hold. Assigning taxonomy to this material is the central computational problem of environmental mycology, and a growing body of work applies deep learning to it. Models trained end to end on barcode sequences report genus- and species-level accuracies above 90% for fungal ITS [22, 16] and for animal COI [4, 7], and independent benchmarking on full-length long-read ITS finds them competitive with established methods for recovering community composition [20]. The barcoding work sits inside a larger literature of pretrained nucleotide models, from early masked-language models of genomic DNA [14] to corpus-scale foundation models [10, 8], which supplies both the architectures and, as we note below, the evaluation practices the barcoding literature has been slower to adopt.

A second strand builds novelty handling into the model rather than treating it as a post-hoc score, either through Bayesian nonparametric priors that admit unobserved taxa at each rank [25] or through deep hierarchical Bayesian formulations [5]; a related line uses DNA as side information for zero-shot recognition of unseen taxa [6]. We do not evaluate these, and our conclusions concern the pretrained fixed-label classifiers currently distributed for fungal ITS. A third bounds what any method can achieve on this marker, and so bounds our own baselines. dnabarcoder [24] showed that similarity cutoffs for fungal identification vary substantially between clades and predicted clade-specific cutoffs accordingly, which is why we treat a single global cutoff as a baseline to be interrogated rather than a standard to be defended; and the accuracy and precision of ITS-based identification have been assessed directly [15], which is part of why we score recovery at family rank throughout rather than treating genus or species as the target.

The reported accuracies are nonetheless difficult to translate into an expectation for environmental data, for two reasons. The first is how test sets are built. Reported figures are typically obtained under random cross-validation in which test sequences are drawn from the same labelled pool, frequently the same genera, as the training data. The extent of that overlap has been quantified: for the largest of the three MycoAI test sets, comprising 363,420 barcodes and the one on which overall performance is usually reported, the overlap analysis reported by Gao and colleagues finds complete overlap with the training set on both measures they report: 363,420 of 363,420 test barcodes are identical to a training barcode and 14,742 of 14,742 test species are present in training, each 100.00% ([7], Table 6). Their two smaller test sets overlap at 86.73% and 6.48% of barcodes, and they describe the two high-overlap sets as measuring performance on in-distribution sequences, naming the 6.48% set as the rigorous test of generalization. Where generalization is tested at all it is tested at the level of unseen species, with the query’s genus and family still represented in training. The regime that characterizes dark matter, a sequence whose entire genus is absent, is not evaluated.

The problem is not new and the surrounding literature has answers to it: models of genomic DNA routinely partition train and test by sequence clustering or by homology, rather than at random, precisely so that a test sequence is not a near-duplicate of a training one [14, 10, 8]. Those partitions transfer imperfectly to environmental barcoding, for reasons worth stating: many species are represented by a single reference sequence, so clustering cannot hold out a within-cluster relative without holding out the taxon entirely, and the marker is hypervariable enough that identity-based clustering separates congeners inconsistently. The stratification we use below is a response to the same concern under those constraints, taking the model’s own label space as the partition rather than a similarity threshold we would have to choose.

The second reason is less often remarked. Reported accuracies are obtained on full-length reference sequences, whereas environmental surveys sequence a primer-bounded subregion. Whether performance on the former predicts performance on the latter has not been established, and nothing in the distributed models or their documentation says it should.

Stated plainly, the question this paper asks is whether the accuracies reported for pretrained fungal ITS classifiers survive the two conditions that define their intended application: a query whose genus the model has never seen, and a query consisting of the ITS2 amplicon alone. Neither condition is exotic. Together they describe most of what an environmental survey submits for classification.

We answer it by benchmarking the two pretrained classifiers against three methods that are fitted or referenced by the user, under conditions designed to be symmetric. Queries are drawn one per genus from the public release the classifiers were trained on, and stratified by whether their genus lies in the classifiers’ own label space, recovered directly from the distributed models. The reference the other methods consult is built to mirror that label space, so a seen-genus query has same-genus references available exactly as the classifiers had same-genus training data, and a novel-genus query has neither. The entire comparison is then run twice, on full-length ITS and on the ITS2 subregion of the identical records, and an ablation on the flanking regions establishes why the two differ.

The question is not untouched. Fujisawa and Imai [12] benchmark conventional and deeplearning methods for insect DNA barcoding under incomplete reference databases and report that detecting species absent from the reference is substantially harder than identifying those present, while finding both classes of method accurate for assignment itself. Our study differs in marker, kingdom and, most consequentially, in comparing amplicon regions, and we reach a stronger conclusion about the classifiers: on our data they lose on assignment as well. Whether that difference reflects the marker, the reference, or the region mismatch we identify below is not settled by either study.

Our results are unfavourable to the classifiers in a specific and, we think, consequential way. They are already outperformed by both classical methods on full-length ITS. On ITS2 they lose between 49.6 and 58.0 percentage points of family accuracy where the classical methods lose between 3.5 and 3.7, and the ablation locates the reason in a dependence on the flanking regions so strong that supplying the flanks of an unrelated phylum causes the transformer to return that phylum’s family for most queries. And on ITS2 neither model’s confidence score retains useful novelty signal, so nothing in the output reports that the query’s genus lies outside the label space; the convolutional model is overconfident there as well, so its names are both mostly incorrect and scored as though they were not. A companion study [18] addresses the complementary question of how abstention can be calibrated once a suitable score exists.

## 2 Methods

Five methods are compared throughout, and it is worth naming the classes they fall into before the details, because the contrast this study draws is not the one a learned-versus-classical framing would suggest. *Best-hit alignment* and *SINTAX* consult a labelled reference at query time and are called the reference-based methods here. *HiTaC* is a hierarchical classifier fitted by the user; *MycoAI-CNN* and *MycoAI-BERT* are classifiers distributed already trained. All three of the latter have fixed label spaces and consult no reference at query time. We avoid opposing “*k*-mer methods” to “classifiers”, since the categories overlap at both ends, and in this study they overlap almost completely: SINTAX classifies, HiTaC is a classifier over 6-mer frequencies, and MycoAI-CNN classifies a 4-mer frequency spectrum (§2.1). The best method at both loci and the worst therefore take input of the same kind, differing in *k* and in what is fitted on top of it.

The axis that matters here is whether a method is fitted or referenced at the locus it is applied to, not which sequence representation it uses.

### 2.1 Classifiers and recovery of their label space

We evaluated the two architectures distributed with MycoAI [22]: a multi-head convolutional network (MycoAI-CNN, 246 MB) and a transformer (MycoAI-BERT, 1.1 GB), both pretrained on the UNITE+INSD dataset as released in 2023 [1], approximately 5.23M sequences after preprocessing [3], spanning 791 families and 3,695 genera, and both obtained from Zenodo record 10904344.

The two differ in how they represent a sequence, and the difference is larger than the shared label space suggests. It also bears directly on the ablation of §3.3, so we set it out here. The convolutional network does not receive the sequence at all. Its input is a normalized frequency spectrum of overlapping 4-mers, a single vector of 625 counts covering the four nucleotides plus a placeholder for ambiguous positions, presented as one channel to a shallow one-dimensional convolutional stack of two layers carrying five and ten filters. That stack holds 320 fitted parameters in total. The model’s remaining 6.6M sit in a fully connected layer and the six per-rank output layers, the largest of which spans 14,742 species, so all but a rounding error of this artefact’s capacity is in naming rather than in reading. The training corpus was encoded this way before fitting. Position is discarded by that encoding: two sequences with the same 4-mer composition in any order are the same input. MycoAI-CNN is therefore a *k*-mer frequency classifier in the same representational family as HiTaC, which fits logistic regressions on 6-mer frequencies, and differs from it in *k*, in what is fitted on top, and in the size of the corpus rather than in kind. The transformer is different: its input is a byte-pair tokenization of the sequence, a vocabulary of learned variable-length units over a bounded number of token positions, and it retains order. These settings are read from the distributed checkpoints themselves, which report their own configuration: MycoAI-CNN declares a KmerSpectrum encoder with *k* = 4, unit stride and normalization in front of a two-layer convolutional stack, and MycoAI-BERT a BytePairEncoder with a vocabulary of 768 units over 256 token positions in front of an eight-layer transformer of model dimension 512. Both models emit the six ranks from separate output heads over separate label encoders, which bears on the coherence diagnostic in §3.2: nothing in the architecture requires the family a model returns to lie inside the order it returns.

A fixed-label classifier can be leakage-controlled only with respect to its own training set, since a genus cannot be withheld from an already-trained softmax head. We therefore recovered each model’s label space directly from its per-rank label encoders rather than reconstructing a training set. The two label spaces were identical at every rank, so a single query set, stratification and reference serve both and the architectures are compared pairwise. Recovering the label space from the distributed model makes the stratification exact and independent of reference release, and the resulting taxon lists are deposited with the harness. They agree exactly, at every one of the six ranks, with the label inventory MycoAI publishes alongside the models, which lists 18 phyla, 70 classes, 231 orders, 791 families, 3,695 genera and 14,742 species; the stratification therefore does not rest on our extraction code alone.

Two properties of the distributed artefacts bear on the ITS2 arm below and are worth stating before the results rather than after them. First, the training corpus consists almost entirely of full-length sequences, which carry the ITS1 spacer and the conserved 5.8S gene that a primer-bounded ITS2 amplicon does not (Figure 1). We describe these throughout as the flanking regions rather than as conserved context, since ITS1 is hypervariable and is itself a standard fungal barcode; what distinguishes them from ITS2 here is position in the training record, not conservation. The corpus is also markedly longer than an ITS2 amplicon: MycoAI preprocessing excludes sequences more than four standard deviations from a mean length of 558.0 bp (SD 126.2), as reported for that dataset by Gao and colleagues [7], against a median of 169 bp for the ITS2 queries. We note the relevant difference is compositional rather than metric, since Section 3.3 shows that restoring length alone does not restore accuracy. Second, neither the MycoAI publication [22] nor the distributed package scopes the models to full-length input, states a minimum query length, or warns on short queries; the mycoai-classify interface accepts primer-bounded ITS2 without error and returns a family score for every sequence submitted. The ITS2 arm is therefore a use the distributed artefact permits and does not flag, which is the sense in which its behaviour there is a property of the released tool and not only of the architecture. Two further operational properties of the package must be worked around before a first prediction is possible, a telemetry call raised on import and a checkpoint recent PyTorch releases decline to unpickle. Neither affects any value reported here, and both are described with their workarounds in Appendix A, since a paper about what practitioners would download should say what downloading it entails.

**Figure 1:**
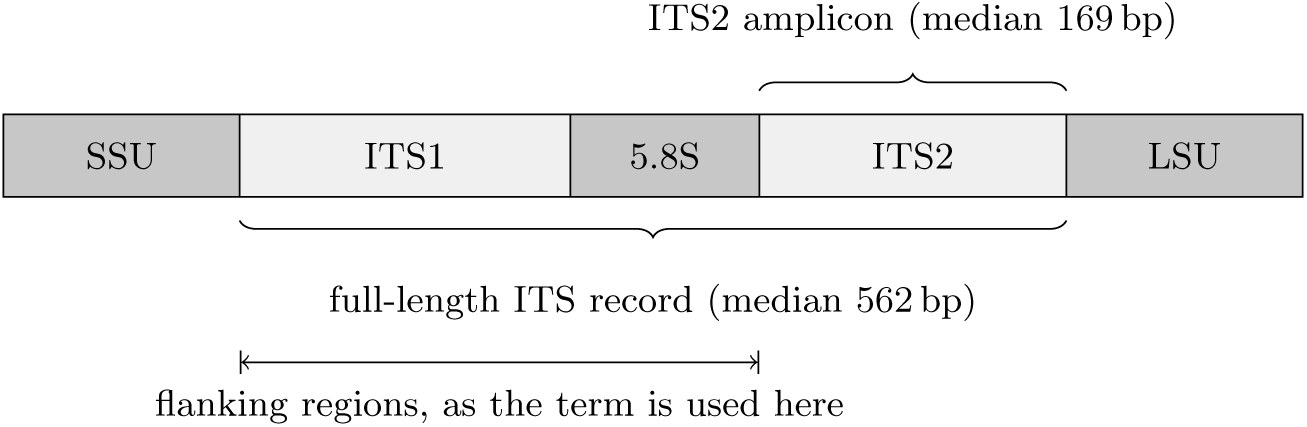
The nuclear ribosomal ITS region and the two loci compared. Reference databases such as UNITE hold the full-length record, which is what the classifiers were trained on and what the first arm supplies. Environmental surveys amplify one spacer, most often ITS2, with primers sited in 5.8S and LSU, so the sequence a survey submits for classification is the ITS2 block alone, which is what the second arm supplies. The two hypervariable spacers are shaded light and the conserved rRNA genes dark; note that ITS1 is a standard fungal barcode in its own right, so “flanking” here denotes position in the training record rather than conservation (§2.1). Drawn to approximate scale.

### 2.2 Query construction and the seen/novel stratification

Queries were drawn from the UNITE+INSD 2024 ITS reference distributed with dnabarcoder [24] (Zenodo 13336328), built from the 21.04.2024 release [2]. That is one release later than the one the classifiers were trained on, which bears on how the seen-genus stratum should be read: a seen-genus query is a sequence whose genus the models learned, but not necessarily a sequence they saw, so seen-genus recovery is a same-genus generalization test rather than a test of recall. The stratification itself is unaffected, being taken from the models’ own label encoders rather than from any release.

A query whose genus lies in the classifiers’ label space is *seen*; one whose genus is absent is *novel-to-model*, shortened to *novel* where no ambiguity arises. This is the distinction the barcoding machine-learning literature draws as *seen* against *unseen* [4, 13]. We keep “novel” because it is the term environmental mycology uses for the same material, and because the stratum is defined here by an already-trained model’s label space rather than by a train/test split we controlled, but the two vocabularies denote the same partition and readers coming from either literature should read them as interchangeable.

Exactly one sequence was taken per genus, eliminating genus-level pseudoreplication so that each genus contributes a single independent test of placement and no genus is over-represented by sequence abundance. This yields 3,330 seen-genus and 1,892 novel-to-model queries. A further 252 genera fell outside the models’ family space and were excluded, correct family placement being impossible by construction. We reconciled those names against GBIF and Catalogue of Life for 195 of the 252, the subset whose records survive ITS extraction. Every one was matched by the nomenclator and exactly one proved to be a synonym, so the excluded set is not appreciably composed of obsolete or superseded names. Thirteen of the 195, 6.7%, nonetheless carry an accepted name whose family lies inside the models’ label space, in almost every case because the nomenclator assigns the genus to a different family than the release does. For those, correct family placement was not impossible by construction after all, and the exclusion is conservative to about that degree. Because the 195 are the excluded genera whose records survived extraction rather than a random draw from the 252, we read 6.7% as indicative of the size of this bias rather than as a measurement of it.

One sequence per genus is a deliberate simplification and it has two costs worth stating here rather than in the limitations alone. It discards intrageneric variation, which for a multicopy region such as ITS is substantial: the same species can carry several divergent and equally valid ITS variants, and a single draw represents none of that spread. And it equalizes genera, not families, so a family represented by many genera contributes more queries than one represented by few, leaving the pooled family-rank accuracies weighted by generic richness. The second is addressable from the deposited per-query tables, and we report every family-rank figure macro-averaged over families alongside the pooled value in Appendix E: macro-averaging lowers every method, by more for the two pretrained models than for the other three at full length, and changes no conclusion. The first would require rebuilding the query set and we have not done it. Because one-per-genus sampling also removes the abundance weighting a random draw from UNITE would carry, absolute recovery under this design is expected to fall below published accuracies obtained on abundance-weighted test sets; the comparisons of interest are between strata, between methods and between loci, all of which share the design.

### 2.3 Paired loci

Because ITS2 is shorter than the full ITS region on which the classifiers were trained, the comparison was run over two paired arms spanning the identical 5,222 records: the full ITS sequences (median query length 562 bp), and the ITSx-extracted [9] ITS2 subregion of those same records (median 169 bp), both distributed with the same release. Restricting the second arm to the record identifiers of the first makes region the only variable, so that any difference is attributable to the amplicon window rather than to taxon sampling.

### 2.4 Reference construction for the reference-based methods

Queries and reference are two disjoint sets. A record enters the reference if and only if its genus lies in the classifiers’ label space and it is not itself one of the queries, capped at twenty sequences per genus; no query sequence appears in any reference at either locus. This gives 56,327 sequences across 3,346 genera for the full-ITS arm and 54,108 for the ITS2 arm, and equalizes the taxonomic knowledge available to all methods. Leakage control is thereby symmetric and automatic. A seen-genus query has same-genus references available, mirroring the classifiers having trained on that genus, while a novel-genus query has none, its genus lying outside the label space and therefore outside the reference by construction.

Disjoint record identifiers are not the same as absent near-duplicates, and the distinction matters for how a high seen-genus score should be read: a query with a nearly identical reference neighbour is being placed by something closer to retrieval than to generalization. The alignment arm measures exactly this as a by-product, its best-hit percentage identity being each query’s identity to its nearest reference neighbour, and the two strata differ sharply on it: median 99.4% for seen-genus queries against 85.5% for novel-genus ones at full length, with 59.3% of the seen stratum having a neighbour at 99% or above and 26.5% an exact match. Removing those near-duplicates costs the reference-based methods and HiTaC around four percentage points of seen-genus recovery and the two pretrained models under one, so part of the leaders’ seen-genus margin is indeed retrieval; the ordering and the size of the ITS2 collapse are unaffected.

Appendix F gives the distributions and the recomputed comparison. To establish that the comparison does not depend on the per-genus cap, the full-ITS reference was rebuilt and rescored at five and fifty sequences per genus, giving references of 16,352 and 113,455 sequences; the ranking of methods is unchanged at every depth, and that analysis is reported in Appendix D.

### 2.5 The five methods

#### Best-hit alignment

Queries were searched with vsearch [21] (usearch_global, best hit, identity floor 0.5, both strands) and assigned the taxonomy of the best hit; best-hit percentage identity served as its score. Of 5,222 queries, 5,220 obtained a hit on full ITS and 5,177 on ITS2; those without were scored as family-incorrect and maximally novel, no reference match meaning no placement.

#### SINTAX

We scored SINTAX [11] as implemented in vsearch, a *k*-mer bootstrap consensus classifier standard in metabarcoding pipelines, which abstains by withholding assignments below a confidence threshold. The identical reference was rewritten with full lineage headers, retaining 55,924 of 56,327 records on full ITS and 53,742 of 54,108 on ITS2, the 0.7% difference being records lacking a complete kingdom-to-genus lineage. Queries were classified with –sintax_cutoff 0.8, the conventional setting, and per-query family-rank bootstrap confidence served as its score.

#### HiTaC

To separate the effect of learned classification from that of the sequence representation, we additionally trained HiTaC [16], a hierarchical classifier that fits a local logistic regression per parent node of the taxonomic tree on 6-mer frequency features. Unlike the MycoAI models, HiTaC is fitted by the user rather than distributed pretrained, so we trained it on the same label-space-mirrored reference the reference-based methods consult. Its label space is therefore that reference’s taxa, novel-genus queries are novel to it by construction, and leakage control is symmetric with the other methods without any need to recover a training taxon list. We fitted it at two reference depths, 16,223 and 55,924 sequences after lineage completion, corresponding to 0.31% and 1.07% of the MycoAI training corpus. A header-format correction its parser requires is described in Appendix A; no family value reported here is affected by it. Its confidence score requires a separately fitted hierarchical filter, which we did not train, so HiTaC contributes no novelty AUROC anywhere below.

#### The pretrained classifiers

Both MycoAI models were run through the distributed mycoai-classify interface with default settings, and the family-rank output served as each model’s score. We call that output a confidence score throughout rather than a probability. It is a softmax over the family label space, and a softmax output is not a calibrated probability unless it has been shown to be one: a returned value of 0.9 does not by itself mean a nine-in-ten chance of being correct. We neither fitted nor evaluated a calibration for these models, and every use we make of the score is ordinal, either as a ranking for AUROC or as a threshold, which calibration is not required for. Whether the scores are in fact calibrated is a separate question, and one worth answering because the interface returns them to users as though they were probabilities; we report reliability diagrams, expected calibration error and Brier scores for both models and for the SINTAX bootstrap in Appendix G, and §3.5 gives the part of that result which bears on the argument.

Training and inference costs were modest for every method and are reported in Appendix B; the largest single cost anywhere in the study is about five minutes of CPU inference for the transformer over the full query set, and the two HiTaC fits took about ten minutes each.

### 2.6 Flank ablation

The ablation is a perturbation-based attribution experiment: the input is modified in controlled ways and the change in output is read as evidence about which part of the input a prediction depends on. It shares that logic with the occlusion and input-ablation analyses used to interpret sequence models generally, and inherits their principal limitation, which is that they establish dependence rather than mechanism (§3.3).

To test directly whether the classifiers depend on the regions flanking ITS2 rather than on ITS2 itself, we constructed seven arms over the same records, all sharing the query identifiers of the full-ITS arm so that a single truth table scores every arm. Each record’s ITS2 was located within its own full ITS by substring match, succeeding for 5,221 of 5,222 records and yielding a median 5*^′^* flank of 360 bp, comprising ITS1 and 5.8S, and a median 3*^′^* flank of 26 bp, an LSU remnant. Region boundaries for the component arms were taken from ITSx [9] (v1.1.3, fungal profiles), which annotated ITS2 for 5,214 records and agreed with the release’s own ITS2 boundary exactly for 5,188 of 5,212 comparable records; median ITS1 is 181 bp and median 5.8S 158 bp, so arm (iv) supplies roughly twice as much donor ITS1 as donor 5.8S and cannot on its own say which the models follow.

The arms are: (i) ITS2 alone; (ii) ITS2 extended on its 5*^′^* side with random ACGT sequence to the training-corpus mean of 558 bp, which adds length without adding information; (iii) ITS2 with its own flanks, which is the full-ITS arm; (iv) ITS2 grafted between the flanks of a donor record drawn from a different phylum, which adds genuine flanking context belonging to the wrong taxon; and three component arms that add back one native region at a time, (v) the record’s own 5.8S followed by its own ITS2, (vi) its own ITS1 followed by its own ITS2, and (vii) its own ITS1 alone with no ITS2 at all. Sixteen phyla were available as donors and no donor shared its query’s phylum. Arms (v) to (vii) use the query’s own sequence throughout rather than a donor’s, so they measure how much of full-length performance each region restores rather than which lineage a prediction follows; the two questions are separate and arm (iv) answers the second.

For arm (iv) each prediction was scored against the donor’s lineage as well as the query’s. Because the query’s ITS2 is present and unaltered while only the flanks belong to the donor, agreement with the donor measures which part of the input the prediction follows, rather than inferring it from an accuracy profile. Arm (i) supplies the chance baseline for donor agreement, no donor sequence being present there.

Arms (ii) and (iv) are constructed inputs that no sequencing workflow would generate. Their purpose is diagnostic, and neither is an estimate of field performance. Arm (ii) in particular confounds two things, since random sequence is both uninformative and potentially misleading, so its result should not be read as a pure length control.

### 2.7 Robustness to the ITS2 delimitation

The ITS2 arm depends on where the ITS2 boundary was drawn, which was done upstream by the release’s authors. The purpose of this arm is narrow: to establish that the ITS2 result below is not an artefact of that one delimitation, by re-extracting ITS2 with an independent implementation and checking that the models behave the same way on both versions. It is a negative control, and the outcome that would support the rest of the paper is no difference between the two.

ITS2 was therefore independently re-extracted from the same full-ITS queries with ITSxRust [19] (v0.2.2, –region its2, bundled fungal HMM profile set, default inclusion *E*-value 10*^−^*^5^, no platform preset), and both extractions were classified on the identical subset of records. Because the queries are ITS-only sequences with a median 3*^′^* LSU remnant of 26 bp, the HMM anchoring required to bound the 3*^′^* end of ITS2 succeeded for 2,637 of 5,222 records, of which 1,754 derived from partial anchor chains; the comparison is therefore restricted to those 2,637 records and its absolute values are not directly comparable with those from the full query set. The corresponding competing interest is declared below.

### 2.8 Scoring and diagnostics

Family-level recovery was scored within each stratum by exact match against the reference taxonomy, pooled over queries. Because all five methods were scored on the identical queries, differences between methods within a stratum were tested with McNemar’s test. That test compares two methods on paired binary outcomes by looking only at the queries where exactly one of the two is correct and asking whether the split between them departs from even; queries both get right, or both get wrong, carry no information about which is better and are discarded. We used the exact binomial form since some discordant counts are small, with Holm adjustment across the forty comparisons, ten pairs of methods in each of the two strata at each of the two loci, all at family rank.

The test addresses variability across queries rather than variability across fits, so it is worth saying why the latter needs no separate treatment here. The two distributed models are fixed artefacts. HiTaC fits a logistic regression per parent node with liblinear in the primal, a configuration in which the solver performs no shuffling and the fit is a deterministic function of the training data; refitting on the same reference reproduces the same classifier, and repeating it under different seeds would measure nothing. What that does not bound is sensitivity to which reference records were drawn, which is a different question and one the depth sweep of Appendix D addresses in part.

Novelty separability of each method’s own score was quantified as the area under the ROC curve separating novel-genus from seen-genus queries, with percentile confidence intervals from 2,000 bootstrap resamples stratified on the two classes. AUROC uses the score only ordinally, so it is well defined for an uncalibrated score, and it measures separation of the two strata rather than correctness of any individual prediction; §3.5 says what follows from that and what does not. As diagnostics distinguishing out-of-domain input from taxonomic failure we additionally report a per-rank accuracy profile from phylum to genus, and the internal coherence of each prediction, meaning whether the predicted family belongs to the predicted order in the reference taxonomy. The seen-genus stratum functions as a positive control throughout: its sequences come from the database the classifiers were trained on and their genera lie in the label space, so low recovery there indicates that the input, not the taxonomy, is out of distribution.

## 3 Results

### 3.1 Classical methods outperform the classifiers on full-length ITS

On full-length ITS, best-hit alignment recovered the correct family for 92.3% of seen-genus and 67.2% of novel-genus queries and SINTAX for 92.0% and 70.3%, against 77.9% and 57.7% for the transformer and 76.5% and 54.3% for the convolutional network (Table 1, Figure 2a,b). Genus-level recovery on seen genera placed the three leading methods well ahead of both distributed classifiers, HiTaC first at 84.8% against 84.4% for SINTAX and 84.3% for alignment, and 67.3% and 71.7% for the transformer and the convolutional network, although the two classifiers exchange places relative to their family ordering. A trained hierarchical classifier fitted to the same reference matched alignment exactly on seen genera at family rank (92.3%), led at genus rank and shared the top of the novel-genus ranking with SINTAX (71.4% against 70.3%, not distinguishable), exceeding best-hit alignment, having been fitted in about ten minutes to 1.07% of the MycoAI training corpus.

**Figure 2:**
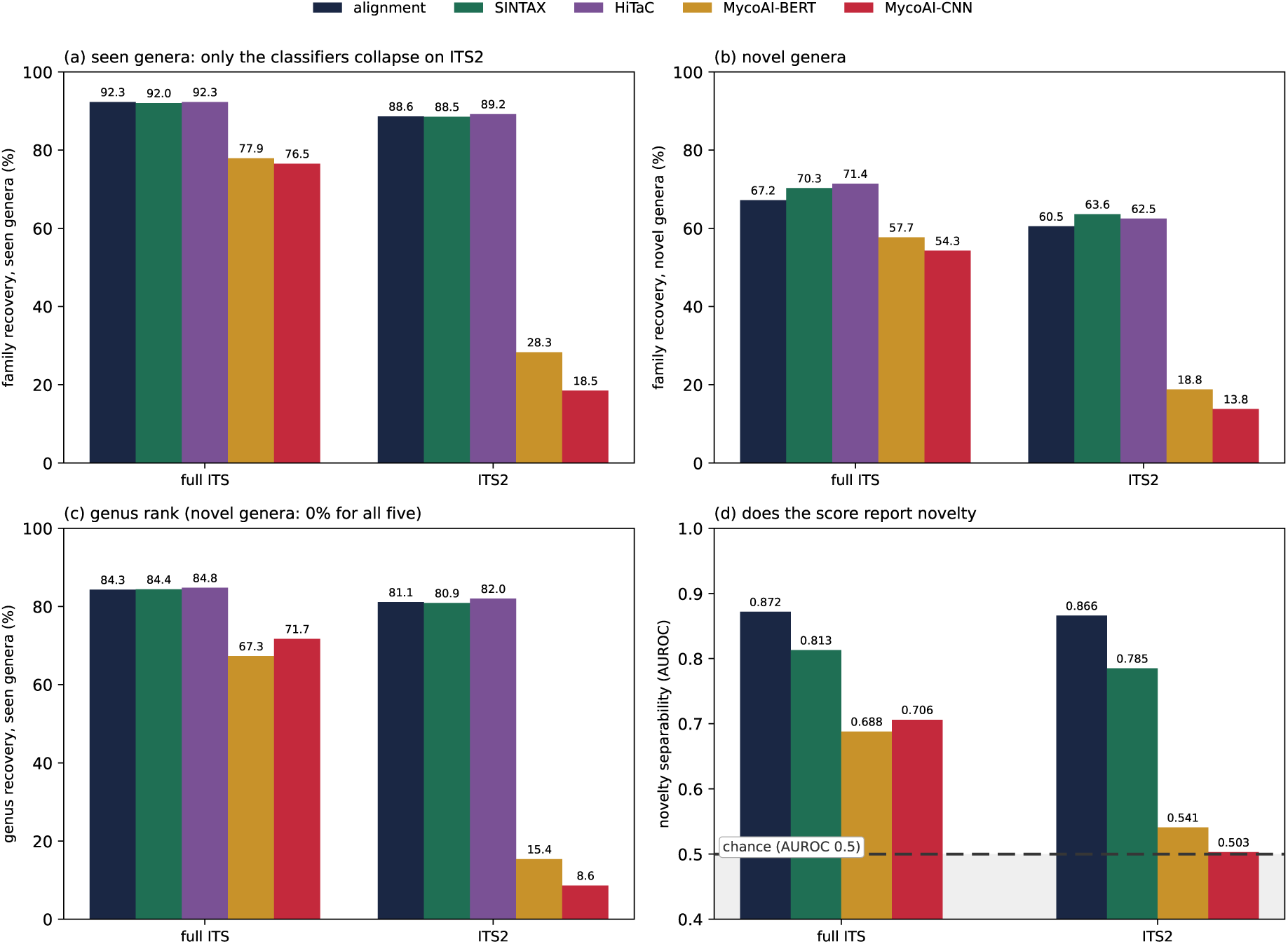
Five methods on 5,222 identical UNITE records, one per genus, stratified by whether a query’s genus lies in the classifiers’ training label space (3,695 genera, recovered from the distributed models) and correspondingly in the reference the reference-based methods consult. Each panel compares full-length ITS with the ITS2 subregion of the same records. (a) Family recovery on seen genera. (b) Family recovery on novel genera. (c) Genus recovery on seen genera; on novel genera it is exactly zero for all five methods at both loci, the correct label being absent from vocabulary and reference alike. (d) Novelty separability of each method’s own score, with chance marked, for the four methods that report one; HiTaC’s requires a filter we did not fit (§2.5). Methods are ordered with best-hit alignment first throughout, as the classical baseline the rest are read against, matching the row order of Table 1; in panel (d) the region below chance is shaded.

**Figure 3:**
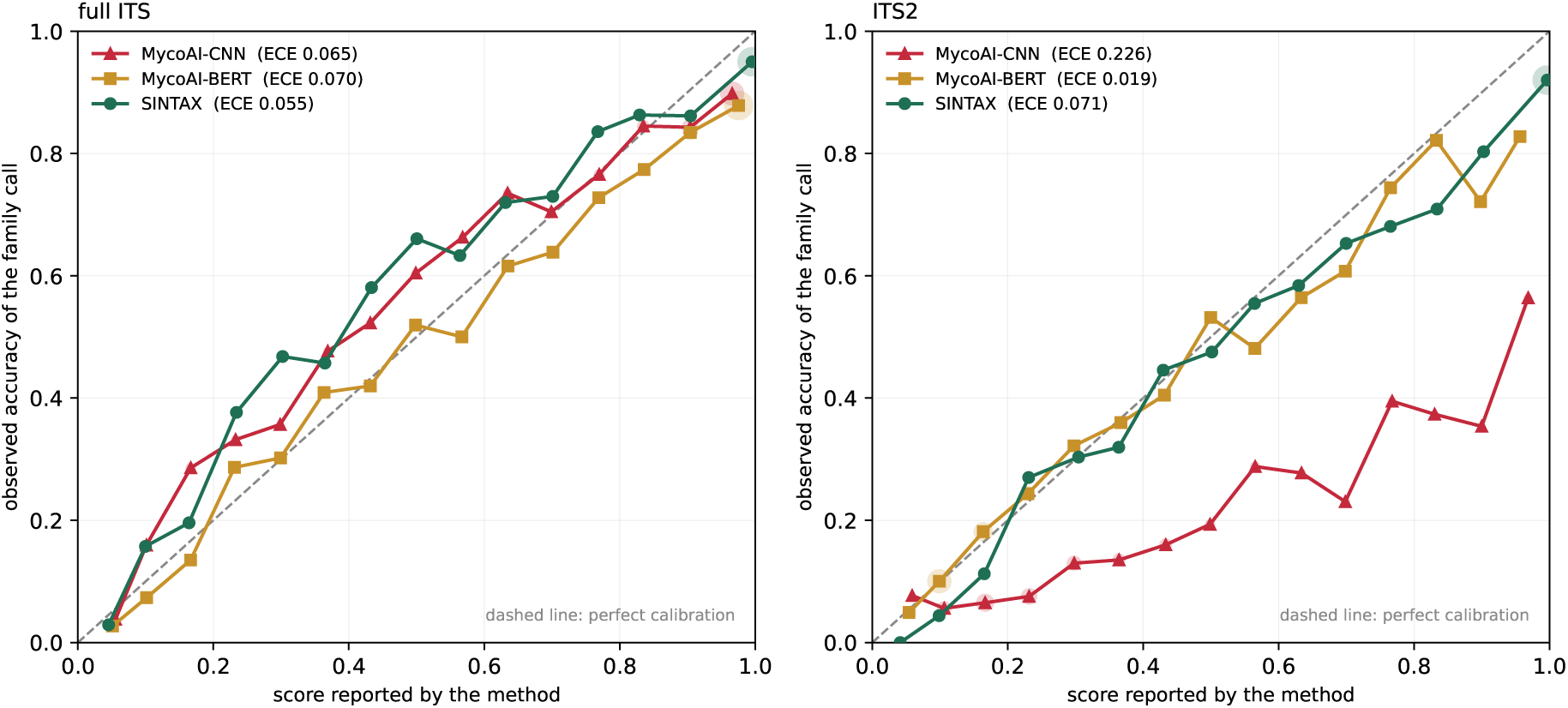
Reliability of the three scores that are offered as confidences, on all 5,222 queries at each locus. Points give observed accuracy of the family call against the mean score in each of fifteen bins, with marker area proportional to the number of queries in the bin and bins holding fewer than 25 queries omitted from the line. A perfectly calibrated score follows the diagonal; points below it are overconfident. On full-length ITS all three track the diagonal closely. On ITS2 the SINTAX bootstrap and MycoAI-BERT continue to do so, while MycoAI-CNN sits well below it across the whole range.

**Table 1:** Five methods on identical queries across both loci. Highest value in each column and locus in bold, which is not the same as a demonstrated lead: on seen genera the three leading methods are not distinguishable at either locus, and genus-rank contrasts were not tested (*§*3.1). Every method is given the resources available at the locus tested: the search methods a reference extracted at that locus, HiTaC a fit at it (*§*3.7), while the two MycoAI models are single distributed artefacts and are the only methods here that cannot be given it.

| locus | method | family<br>seen genus | family<br>novel genus | genus<br>seen genus | novelty<br>AUROC |
| --- | --- | --- | --- | --- | --- |
| full ITS | alignment | **92.3%** | 67.2% | 84.3% | **0.872** |
|  | SINTAX | 92.0% | 70.3% | 84.4% | 0.813 |
|  | HiTaC | **92.3%** | **71.4%** | **84.8%** | n/a |
|  | MycoAI-BERT | 77.9% | 57.7% | 67.3% | 0.688 |
|  | MycoAI-CNN | 76.5% | 54.3% | 71.7% | 0.706 |
| ITS2 | alignment | 88.6% | 60.5% | 81.1% | **0.866** |
|  | SINTAX | 88.5% | **63.6%** | 80.9% | 0.785 |
|  | HiTaC | **89.2%** | 62.5% | **82.0%** | n/a |
|  | MycoAI-BERT | 28.3% | 18.8% | 15.4% | 0.541 |
|  | MycoAI-CNN | 18.5% | 13.8% | 8.6% | 0.503 |

The comparison is therefore not between learned and classical methods, and framing it that way would misdescribe the result. Learned classification works well here: HiTaC is a trained, fixed-label classifier and it placed novel genera better than either reference-based method. What underperforms is the pair of distributed MycoAI models, which are exceeded on full-length ITS by best-hit alignment, by SINTAX, and by a hierarchical logistic regression on *k*-mer counts fitted to one percent of their training data. That is a narrower claim than the one a learnedversus-classical framing would support, and a harder one to attribute to the choice of baseline.

Thirty of the forty pairwise contrasts survive McNemar’s test after Holm adjustment. Nine of the ten that do not involve only the three leading methods, which on seen genera are mutually indistinguishable at both loci: 92.3%, 92.0% and 92.3% on full ITS and 88.6%, 88.5% and 89.2% on ITS2, every pairwise adjusted *p* equal to 1.00, with alignment against HiTaC splitting the discordant pairs exactly 75 to 75 at full length. On novel genera the result is a partial ordering rather than a tie. At full length SINTAX and HiTaC are indistinguishable (70.3% against 71.4%, adjusted *p* = 1.00) and both exceed alignment (1.9 *×* 10*^−^*^6^ and 1.8 *×* 10*^−^*^5^); on ITS2 SINTAX exceeds alignment (9.7 *×* 10*^−^*^7^) while HiTaC is distinguishable from neither (1.00 and 0.34). Every contrast involving a MycoAI model is significant in every cell at both loci. The tenth undistinguished contrast is the transformer against the convolutional network on seen genera at full length, 77.9% against 76.5%, adjusted *p* = 0.071.

Two things follow. Where a same-genus reference is available, no difference was detected among the three leading methods at either locus. That is not a demonstration that they are equivalent: the study was not designed or powered as an equivalence test and places no bound on how large a true difference could be while still going undetected on 5,222 queries, so we neither interpret the small gaps between them nor describe the methods as substitutable. Where a same-genus reference is not available, the two that aggregate evidence across several relatives, SINTAX by bootstrapping *k*-mer subsamples to a consensus and HiTaC by fitting a classifier over *k*-mer frequencies, both exceed best-hit alignment, which commits to a single neighbour. That distinction matters most precisely when no same-genus relative exists, and it is the one place in the study where the reference-based and fitted approaches separate in favour of the latter.

Degradation from seen to novel genera was of similar magnitude for every method, 21.7, 25.1, 20.9, 20.2 and 22.1 percentage points for SINTAX, alignment, HiTaC, the transformer and the convolutional model, so the MycoAI models’ disadvantage is close to a constant offset rather than something specific to novel taxa. Capacity helped, but not usefully: the transformer exceeded the convolutional model in all four combinations of locus and stratum, by 1.4, 3.4, 9.8 and 5.0 percentage points, significant after adjustment in three of them, the exception being the 1.4-point gap on seen genera at full length (adjusted *p* = 0.071). It did so while being about 140 times slower at inference, while remaining worse at genus rank on seen genera, and while closing none of the gap to the other three methods, whose advantage over it remains 14 points on full ITS and 60 on ITS2. The two models differ about fourfold in total parameters, which understates the difference in the part that reads the sequence: the transformer’s encoder holds 17.6M parameters against 320 in the convolutional stack, the rest of the convolutional model sitting in output layers whose size is set by the label space rather than by the architecture (*§*2.1). Read that way the comparison is less between two capacities than between a learned encoder and almost none, and the transformer’s margin over a 320-parameter feature extractor is 1.4 points at full length.

Three checks on how these figures were computed leave the picture unchanged, and are reported in full in the appendices. The ranking is not an artefact of how deep a reference the reference-based methods were given: repeating the full-ITS alignment arm at five and fifty sequences per genus leaves alignment ahead of both distributed models in both strata at every depth tested (Appendix D). It is not an artefact of weighting families by their generic richness: macro-averaging family recovery over families lowers every method, the two pretrained models most of all at full length (*−*9.2 and *−*10.1 points against *−*3.0 to *−*4.0 for the other three), and preserves the separation between them (Appendix E). And it is not driven by queries that had a near-identical reference neighbour, though those are common in the seen stratum: discarding every query whose best reference hit reaches 99% identity, which removes three-fifths of that stratum, costs alignment, SINTAX and HiTaC between 3.8 and 4.5 points of seen-genus recovery at full length and the two pretrained models under one point, leaving a lead of ten points rather than fifteen (Appendix F). The last of these is a real qualification and we state it as one: some of the leaders’ seen-genus margin is retrieval of a close relative rather than generalization. None of it touches the ITS2 result, where the same exclusion moves every method by less than two points and the classifiers remain sixty points behind.

### 3.2 On the amplicon surveys actually sequence, only the classifiers collapse

Environmental fungal metabarcoding is dominated by primer-bounded ITS2 amplicons, so we repeated the comparison on the ITS2 subregion of the identical records with a reference built by the same membership rule. The methods separate decisively (Table 1, Figure 2). Seen-genus family recovery fell by 3.7 percentage points for alignment, 3.5 for SINTAX and 3.1 for HiTaC refitted at the locus, but by 49.6 for the transformer and 58.0 for the convolutional network, reaching 28.3% and 18.5%. Novel-genus recovery behaved the same way, falling 6.7 points for both reference-based methods against 38.9 and 40.5 for the classifiers. Genus recovery on seen genera fell from 84.3% to 81.1% for alignment but from 71.7% to 8.6% for the convolutional network.

Because the records, the stratification and the reference construction rule are common to every method, this is not a property of the shorter amplicon. ITS2 carries sufficient information for all three methods that can be given it to retain 96.2%, 96.0% and 96.6% of their full-length family accuracy on seen genera, and between 88% and 91% on novel genera; what fails is the two distributed models’ ability to use it. The per-rank profile locates the failure at the input. On ITS2, convolutional accuracy decayed monotonically from phylum (77.3%, barely above the accuracy of always naming the most abundant fungal phylum) through class (52.9%) and order (30.4%) to family (18.5%), with only 49.6% of predictions internally coherent in the sense that the predicted family belongs to the predicted order. The transformer behaved likewise (76.5%, 60.7%, 41.4%, 28.3%, coherence 55.0%). The corresponding full-ITS figures are 98.8%, 95.3%, 87.5%, 76.5% with coherence 90.5%, and 98.2%, 95.8%, 89.3%, 77.9% with coherence 92.2%. Given an input outside their training domain, these models did not degrade gracefully towards the correct higher ranks; they produced lineages whose own ranks contradict one another.

Two readings of that incoherence are available and they are not exclusive. The first is the one the per-rank profile invites, that an input outside the training domain leaves the prediction unconstrained at every rank at once. The second is architectural: the MycoAI models emit each rank from a separate head over a separate label encoder (§2.1), so nothing in the model or in the distributed inference path requires the family returned to lie within the order returned, and incoherent output is available to these models whenever the per-rank outputs disagree, whatever the input. That is a design choice, and the codebase records it as one. MycoAI implements five output heads, several of which tie the ranks together, either by conditioning each rank on the adjacent one or by deriving parent ranks from the child prediction through the taxonomy. Both distributed checkpoints use MultiHead, the head that predicts the six ranks independently, and the classification interface exposes no argument by which a user could select another: it accepts an input file, an output path, a model path, a confidence flag and a device. The coherence figures above are therefore a property of which head was released as much as of the input the models were given. The two readings predict the same thing here and these data cannot separate them. What distinguishes them is the remedy: the second would be largely repaired by constrained decoding, which would convert incoherent lineages into coherent wrong ones without improving family recovery, and the first would not. We therefore report coherence as a diagnostic of the input while noting that it is partly a property of the models’ output layer, and we do not rest the flank argument of §3.3 on it.

Query length within the ITS2 arm does not explain the effect and the two models do not even agree on its direction: family recovery across length quartiles rose then plateaued for the convolutional model (11.2, 21.7, 18.3, 22.6 per cent) and declined for the transformer (33.0, 34.1, 27.6, 18.9). No coherent account of the collapse follows from length in either direction, and the ablation below shows why: length is not the operative variable.

### 3.3 The classifiers read the flanking regions rather than the barcode

Section 3.2 locates the failure in the input without establishing which property of the input is responsible, and the training-corpus composition noted in Section 2.1 suggests the missing flanking regions. The ablation tests that directly and supports a stronger conclusion (Table 2). It is a perturbation-based attribution experiment in the sense of §2.6: each arm changes one property of the input and the change in output is the evidence. Read that way, an arm answers the question of what a prediction depends on, and does not by itself localize the dependence to particular positions or identify the mechanism behind it; §6 sets out the position-resolved analyses that would.

**Table 2:**
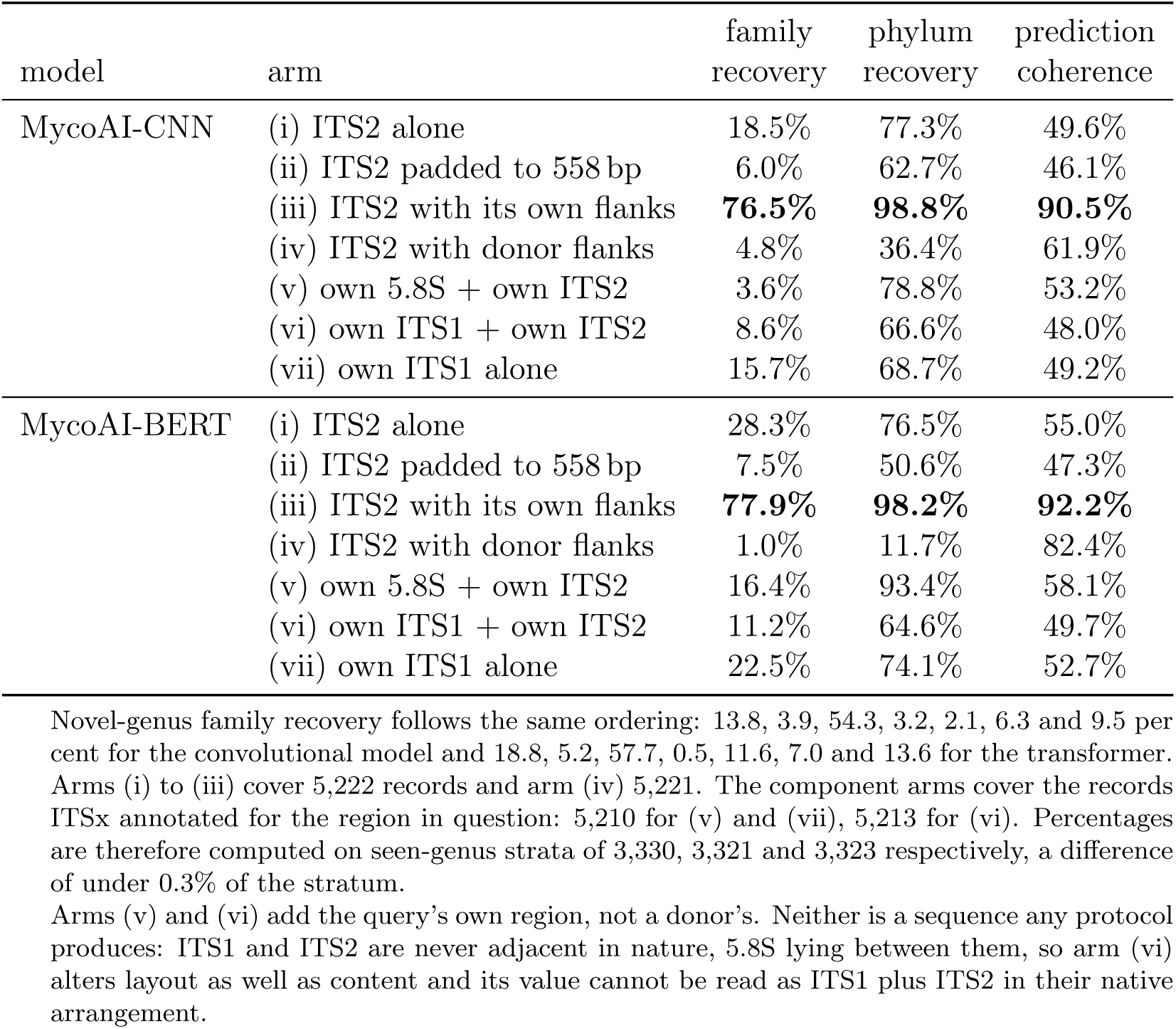
Flank ablation, seen-genus stratum. Every arm covers the same records, 5,222 for arms (i) to (iii) and 5,221 for arm (iv), one record having no locatable flanks, and except where padding is added contains the same unaltered ITS2 sequences. Coherence is the proportion of predictions whose predicted family belongs to the predicted order.

| model | arm | family<br>recovery | phylum<br>recovery | prediction<br>coherence |
| --- | --- | --- | --- | --- |
| MycoAI-CNN | (i) ITS2 alone | 18.5% | 77.3% | 49.6% |
|  | (ii) ITS2 padded to 558 bp | 6.0% | 62.7% | 46.1% |
|  | (iii) ITS2 with its own flanks | **76.5%** | **98.8%** | **90.5%** |
|  | (iv) ITS2 with donor flanks | 4.8% | 36.4% | 61.9% |
|  | (v) own 5.8S + own ITS2 | 3.6% | 78.8% | 53.2% |
|  | (vi) own ITS1 + own ITS2 | 8.6% | 66.6% | 48.0% |
|  | (vii) own ITS1 alone | 15.7% | 68.7% | 49.2% |
| MycoAI-BERT | (i) ITS2 alone | 28.3% | 76.5% | 55.0% |
|  | (ii) ITS2 padded to 558 bp | 7.5% | 50.6% | 47.3% |
|  | (iii) ITS2 with its own flanks | **77.9%** | **98.2%** | **92.2%** |
|  | (iv) ITS2 with donor flanks | 1.0% | 11.7% | 82.4% |
|  | (v) own 5.8S + own ITS2 | 16.4% | 93.4% | 58.1% |
|  | (vi) own ITS1 + own ITS2 | 11.2% | 64.6% | 49.7% |
|  | (vii) own ITS1 alone | 22.5% | 74.1% | 52.7% |

Novel-genus family recovery follows the same ordering: 13.8, 3.9, 54.3, 3.2, 2.1, 6.3 and 9.5 per cent for the convolutional model and 18.8, 5.2, 57.7, 0.5, 11.6, 7.0 and 13.6 for the transformer. Arms (i) to (iii) cover 5,222 records and arm (iv) 5,221. The component arms cover the records ITSx annotated for the region in question: 5,210 for (v) and (vii), 5,213 for (vi). Percentages are therefore computed on seen-genus strata of 3,330, 3,321 and 3,323 respectively, a difference of under 0.3% of the stratum.
Arms (v) and (vi) add the query’s own region, not a donor’s. Neither is a sequence any protocol produces: ITS1 and ITS2 are never adjacent in nature, 5.8S lying between them, so arm (vi) alters layout as well as content and its value cannot be read as ITS1 plus ITS2 in their native arrangement.

Extending ITS2 with uninformative sequence to the training-corpus mean length made matters worse rather than better, reducing seen-genus family recovery from 18.5% to 6.0% in the convolutional model and from 28.3% to 7.5% in the transformer. Length is therefore not the operative variable, which also accounts for the inconsistent length gradients reported above.

Supplying genuine flanking context from the wrong taxon was worse still, reducing family recovery to 4.8% and 1.0% and phylum recovery to 36.4% and 11.7%, in both cases below the values obtained from ITS2 with no flanks at all. A model merely lacking context would be expected to perform no worse when context is restored, whatever its provenance. These models perform much worse, which indicates that the flanks are not supplementary but decisive. Prediction coherence moves in the opposite direction to accuracy, rising to 61.9% and 82.4% against 49.6% and 55.0% on ITS2 alone: the outputs become internally more consistent as they become less correct, which is the signature of a model confidently reporting a lineage, though not the query’s.

Scoring arm (iv) against the donor’s lineage identifies whose (Table 3). Agreement with the donor rises from between 0.1 and 0.7 per cent, where no donor sequence is present, to roughly a third at every rank for the convolutional model and to between 37 and 72 per cent for the transformer. The effect is markedly stronger in the transformer, and the two models differ in kind at phylum rank, where the convolutional model divides evenly between query and donor (37.8% each) while the transformer follows the donor almost seven times more often than the query. At family rank, the rank this benchmark concerns, both follow the donor predominantly. At family rank the transformer returns the donor’s family for 63.7% of queries and the query’s own for 0.8%, a ratio of roughly eighty to one, on sequences whose ITS2 is present and correct. Set against its 77.9% family recovery on native full-length input, it recovers the donor’s family at 82% of the rate at which it recovers the correct one when unperturbed: arm (iv) does not present to it as a damaged sequence but as an intact sequence of the donor organism.

**Table 3:** Whose taxonomy does the prediction follow? Agreement of arm (iv) predictions with the query’s own lineage and with the donor’s, pooled across strata (*n* = 5,221). The query’s ITS2 is present and unaltered; only the flanks belong to the donor, which is always from a different phylum.

| rank | MycoAI-CNN |  | MycoAI-BERT |  | baseline, arm (i) |  |
| --- | --- | --- | --- | --- | --- | --- |
|  | query | donor | query | donor | query | donor |
| phylum | 37.8% | 37.8% | 10.8% | **71.7%** | 78.8% | 0.7% |
| class | 20.8% | 36.3% | 5.4% | **71.3%** | 51.2% | 0.4% |
| order | 10.3% | 38.2% | 2.4% | **68.5%** | 28.8% | 0.3% |
| family | 4.2% | **34.4%** | 0.8% | **63.7%** | 16.8% | 0.1% |
| genus | 0.9% | 26.8% | 0.3% | 36.9% | 5.5% | 0.1% |

Comparing the two arms rules out the obvious alternative reading. In arm (ii) the added sequence is likewise the majority of the input but carries no taxonomic signal, and the result is degradation without misdirection. In arm (iv) the added sequence carries genuine but incorrect signal, and the prediction follows it. The effect is therefore not dilution by input proportion but the models weighting information in the flanking regions above information in ITS2. Which flanking component carries that weight is not resolved here, since arm (iv) transfers ITS1 and 5.8S together (§2.6); ITS1 is the larger of the two and is itself a barcode, so the donor’s ITS1 is the more likely vehicle, but we have not separated them.

The two models differ in how strongly they follow the donor: measured as donor-phylum agreement relative to each model’s own phylum recovery on native input, the convolutional model stands at 38% and the transformer at 73%. We do not read this as a capacity effect. The two models differ in architecture, and specifically in whether their input has a position axis at all (§2.1), which is enough to account for the direction of the difference. The transformer receives an ordered token sequence and can attend to where a *k*-mer occurs; the convolutional network receives a 4-mer frequency spectrum, in which position has already been discarded, so it cannot distinguish a flanking *k*-mer from an ITS2 one and can only respond to the composition of the whole. Arm (iv) is a mixture in that composition: the median record contributes about 386 bp of donor flank against 169 bp of query ITS2, so roughly seven of every ten 4-mers reaching the convolutional model come from the donor. A model that can only average should be pulled part of the way towards the donor rather than switch to it, which is what the even division at phylum rank looks like (37.8% each); a model that retains order can key on the flanking positions and switch, which is what a near-sevenfold preference for the donor looks like. We offer this as the reading the architectures make available rather than as something the ablation tests, since no arm here manipulates order while holding composition fixed. What the comparison supports without it is narrower and is reinforced by §3.7: the more of the input a model is able to exploit, the more it loses when part of that input is withheld.

Restoring one native region at a time asks a different question, and its answer constrains the mechanism further (Table 2, arms (v) to (vii)). Neither component recovers full-length performance and both are worse than ITS2 alone. Adding the record’s own 5.8S to its own ITS2 takes seen-genus family recovery from 18.5% to 3.6% in the convolutional model and from 28.3% to 16.4% in the transformer; adding its own ITS1 gives 8.6% and 11.2%. ITS1 presented alone, with no ITS2 at all, recovers 15.7% and 22.5%, which is no better than ITS2 alone. So these are not ITS1 classifiers that tolerate ITS2 input, and no single flanking region carries the signal the full record supplies.

What the models require is the complete assembly rather than any particular region within it. That is consistent with arm (iv) rather than in tension with it: given something with the shape of a whole record the prediction follows the flanking positions, even to another phylum’s family, while given a fragment it does poorly whatever the fragment contains. Following the flanks and classifying from the flanks are different properties, and only the first is demonstrated here. It also explains why arm (ii) behaves as it does, since random padding and a genuine but partial reconstruction both leave an input that resembles no training record. For the convolutional model the padding arm has a second and simpler reading: adding 389 bp of random sequence to a 169 bp amplicon drives the 4-mer spectrum it classifies most of the way towards the uniform composition of random DNA, which is not a fungal spectrum of any kind.

One dissociation within this is worth reporting separately, because it is the only place where adding context helps anything. In the transformer, appending 5.8S raises phylum recovery from 76.5% to 93.4%, close to the 98.2% it reaches on native full-length input, while family recovery falls from 28.3% to 16.4%. The conserved region restores deep placement and costs shallow discrimination at the same time. The convolutional model shows no such effect, moving from 77.3% to 78.8% at phylum, and we read the difference as consistent with the transformer being able to use where the conserved block sits, which the convolutional model’s order-free input does not let it do (*§*3.3); this is an interpretation and not a measurement. For a practitioner the consequence is narrow but real: a partial reconstruction can make phylum-level output look healthier while making the family call worse, so agreement at deep ranks is not evidence that the input is in distribution.

It is worth separating this account from a weaker one it is easily confused with. Any ITS2 query is, for a model trained on full-length records, an input whose format matches nothing in training, and a general out-of-distribution effect would account for the loss of accuracy on arm (i) with no appeal to the flanking regions at all. The two accounts are nested rather than rival, and the ablation was built to separate them. Format novelty predicts degradation and incoherence; it does not predict which lineage a degraded prediction should name. Arm (iv) restores the format, presenting a full-length-shaped record whose ITS1, 5.8S and ITS2 are all genuine fungal sequence, and accuracy falls further while donor agreement rises to 63.7% at family rank in the transformer, four-fifths of the rate at which that model recovers the correct family from native input. A model merely disoriented by an unfamiliar input format would not name the donor’s family almost as reliably as it names the right one. Arm (ii) makes the complementary point from the other side, restoring the length statistic without restoring genuine flanks and producing degradation with no misdirection at all. What the ablation establishes is therefore that among the ways an ITS2 query differs from a training record, absence of the flanking regions is the one the prediction tracks. It does not establish that an ITS2 input is otherwise in distribution, and §3.5 returns to that point, since it governs how the novelty scores should be read. Quantifying the residue, by embedding full-length records alongside progressively truncated versions of themselves and measuring how far the pair separates, requires access to the models’ internal representations rather than their output and is set out among the experiments of §6.

This is a property of how these models exploit the barcode rather than of ITS2 specifically, and on that reading the same failure would be expected for any protocol that sequences a subregion without its flanking regions.

### 3.4 The collapse does not depend on how ITS2 was delimited

Because the ITS2 arm inherits a boundary drawn by the release’s authors, ITS2 was independently re-extracted from the same queries with a separate implementation and both versions classified on the identical 2,637 records for which re-extraction succeeded (§2.7). The two delimitations proved close, 85.1% of records identical and 99.7% identical or nested, and neither model moved by more than 0.4 percentage points of family recovery between them. The collapse is therefore not an artefact of where this particular boundary was drawn. Because the two extractions are near-identical in sequence, the check establishes independence of implementation rather than of methodology, and does not exclude the possibility that a substantially different ITS2 window would behave differently. Full values, the representativeness of the subset, and the observation that the more flank-dependent model is the less boundary-sensitive are given in Appendix C.

### 3.5 On ITS2 the classifiers’ confidence loses its novelty signal

The more consequential half of the result concerns what the models report about themselves. Treating each method’s own score as a novelty signal, alignment identity separated novel from known genera with AUROC 0.872 on full ITS and 0.866 on ITS2, and the SINTAX bootstrap with 0.813 and 0.785: both degraded slightly and remained strongly informative. The classifiers’ family-rank confidence score fell from 0.688 to 0.541 for the transformer and from 0.706 to 0.503 for the convolutional network (Figure 2d). The convolutional model’s ITS2 value is indistinguishable from chance (95% CI 0.487–0.518) and the transformer’s exceeds it only marginally (0.525–0.557), a distinction that does not survive contact with use: a conformal novelty flag calibrated on these same two scores at a 5% false-novelty rate detects 6.1% of genuinely novel genera for the convolutional model and 5.2% for the transformer while firing on 4.6% and 4.2% of known ones, against 42.7% detection at the same error rate for alignment identity on the same amplicon [18].

Two things this measurement does not say should be stated beside it, because the number invites both readings. First, every ITS2 query reaches these models in a format their training corpus does not contain, so the AUROC reported here is the separation of novel from seen genera *within* that condition (Table 4); it is not a measure of whether the models can recognize that an ITS2 input is unlike their training data at all, which their output does not support asking. Second, detecting a novel genus and knowing whether a prediction is correct are different questions, and on ITS2 these models answer them differently enough that the two must be kept apart.

**Table 4:** What each cell of the design means for a model trained on full-length ITS. The novelty AUROC reported in this section compares the two columns within a row. It does not compare the rows, and no measurement here asks whether a model can tell which row it has been given.

| input given to the model | seen genus | novel genus |
| --- | --- | --- |
| full-length ITS | near the training distribution | taxonomically novel |
| ITS2 | novel in input format | novel in format and in taxonomy |

Taking the second directly, we asked whether each score is calibrated as a probability that the family returned is correct (Appendix G). The two models diverge, and only one of them behaves as the summary above would suggest. MycoAI-CNN is substantially overconfident on ITS2: it reports a mean score of 0.394 while placing 16.8% of queries correctly, an expected calibration error of 0.226 against 0.065 for the same model at full length, and its most confident calls are not rescued by thresholding, the queries it scores above 0.9 being correct 49.8% of the time. MycoAI-BERT is not overconfident on ITS2 at all. Its mean score is 0.252 against 24.9% accuracy and its expected calibration error is 0.019, the lowest of any method, locus or stratum we measured. Its ITS2 scores are simply low: the median is 0.171, and thresholding at 0.8 retains 3.3% of queries, which are then correct 78.7% of the time.

That divergence sharpens rather than softens the practical conclusion, and it changes how we state it. For the convolutional model the score is misleading about correctness and at chance for novelty, so an ITS2 run yields confident names, mostly wrong, with nothing in the output to indicate it. For the transformer the score remains a usable indicator of whether its own call is right, and yet is nearly uninformative about whether the genus is novel (AUROC 0.541); a practitioner who trusts it is therefore left choosing between accepting predictions that are wrong three times in four and discarding some ninety-seven per cent of the run. Calibration and discrimination come apart here as cleanly as they can: a score can be well calibrated and still carry almost no information, since one that reports a quarter and is right a quarter of the time is calibrated by construction. What neither model provides on this amplicon is the thing the application needs, a score that falls when the query’s genus is absent from the label space. This is the basis of our recommendation that reported accuracies specify the amplicon region of the evaluation and be accompanied by a same-query classical baseline.

### 3.6 No method can name a novel genus; they differ in whether they say so

At genus rank all five methods scored exactly 0.0% across all 1,892 novel-genus queries, at both loci. The result is structural rather than empirical and we do not dwell on it: the correct genus is absent from the classifiers’ output spaces and from the reference the other methods consult, so the task as posed puts the right answer out of reach for every method. No increase in training data or model capacity alters this for a fixed-label model, and no reference supplies a name it does not contain.

What separates the methods is disclosure. Alignment reports a best-hit identity that falls when no close relative exists; SINTAX reports a bootstrap confidence that does the same and, by convention, withholds assignments below a threshold. Both therefore emit a quantity from which abstention can be constructed, and on ITS2 both retained that property. The classifiers report a softmax score over a fixed vocabulary which, on ITS2, was uninformative about novelty. Forced prediction from a closed set is thus not by itself the problem, since it applies to every method here. The problem is forced prediction without a score that degrades informatively, and the classifiers on ITS2 are the case where that combination bites. HiTaC is absent from this particular comparison for a reason unrelated to its performance: its novelty score comes from a hierarchical confidence filter fitted separately from the classifier itself, which we did not train (§2.5), so it has no score to assess rather than a poor one.

### 3.7 Training at the amplicon locus removes the deficit

Table 1 gives every method the resources available at the locus being tested, which for HiTaC means a separate fit at each. Comparing those two fits against both query sets isolates what the locus contributes (Table 6). Fitted and tested on ITS2, HiTaC recovered 89.2% of seen-genus and 62.5% of novel-genus families, against 47.4% and 30.8% for the same classifier fitted on full-length ITS and given the identical ITS2 queries. On-locus training returns 41.8 of the 44.9 percentage points lost on seen genera, and the recovered value sits with the search methods rather than beneath them: 89.2% against 88.6% for best-hit alignment and 88.5% for SINTAX, retaining 96.6% of its own full-length accuracy against their 96.2% and 96.0%, with 82.0% genus recovery against their 81.1% and 80.9%.

Two things follow, and the second is what bears on the MycoAI models. First, ITS2 is not the limitation. A hierarchical logistic regression over 6-mer frequencies, fitted in ten minutes to 1.07% of the MycoAI training corpus, places primer-bounded ITS2 amplicons about as well as alignment does, on the amplicon where the distributed models place 28.3% and 18.5%. Neither the shorter window, nor *k*-mer features, nor learned classification is what fails there. Second, the deficit is a mismatch between training and test locus rather than a property of the shorter sequence, which the reverse arm establishes: fitted on ITS2 and given full-length queries, the same classifier fell to 67.7% and 40.1%. Both diagonals are high and both off-diagonals low, so no account in terms of ITS2 carrying less usable information survives the comparison.

The two off-diagonals are not symmetric. Withholding the flanking regions from a classifier fitted with them cost 44.9 points; supplying them to one fitted without them cost 21.5. We report the asymmetry without resting anything on it, and note that it is not the comparison arms (ii) and (iv) of *§*3.3 make: both off-diagonal inputs here are the query’s own sequence, whereas arm (iv) substitutes another phylum’s flanks. What it says is only that a locus-mismatched *k*-mer profile is damaged more by truncation than by extension.

Fitting the full-ITS model at two reference depths adds a second, smaller finding about what governs off-locus robustness (Table 5). Increasing the reference from 16,223 to 55,924 sequences raised full-ITS recovery, from 90.4% to 92.3% on seen genera, and halved ITS2 recovery, from 69.5% to 47.4%; retention fell from 77% to 51%. More training data made the classifier better at the locus it was trained on and substantially worse away from it. Read alongside the locus grid, the reading is straightforward: depth buys features specific to the training locus, so more is lost when the locus changes, and nothing is lost when it does not. The same direction appears in the alignment depth sweep of Appendix D, where deeper references improved placement of known taxa and slightly impaired placement of novel ones. In both cases a larger reference is not uniformly better.

**Table 5:** HiTaC family recovery at two reference depths, on the same queries. Retention is ITS2 recovery as a percentage of full-ITS recovery.

| reference | seen genus |  | novel genus |  | retention |  |
| --- | --- | --- | --- | --- | --- | --- |
|  | full ITS | ITS2 | full ITS | ITS2 | seen | novel |
| 16,223 sequences | 90.4% | 69.5% | 70.5% | 48.5% | 77% | 69% |
| 55,924 sequences | 92.3% | 47.4% | 71.4% | 30.8% | 51% | 43% |

**Table 6:** HiTaC family recovery under every combination of training locus and test locus, on the same queries and at fixed reference membership. Diagonal cells match training and test locus; off-diagonal cells are the mismatch.

| trained on | reference | tested on full ITS |  | tested on ITS2 |  |
| --- | --- | --- | --- | --- | --- |
|  |  | seen | novel | seen | novel |
| full ITS | 55,924 | **92.3%** | **71.4%** | 47.4% | 30.8% |
| ITS2 | 53,742 | 67.7% | 40.1% | **89.2%** | **62.5%** |

Together these results qualify the mechanism proposed in *§*3.3 without displacing it. The flank ablation shows directly that the MycoAI models weight ITS1 and 5.8S above ITS2, and that finding rests on interventions on the input rather than on any comparison across methods. What the present section adds is that the resulting deficit is specific to the mismatch and not intrinsic to the amplicon, since it disappears when a classifier of the same family is fitted at the locus, and that susceptibility to it grows with how much of the training record a model exploits rather than being fixed by architecture. The grid also suggests an experiment we did not run. A single classifier fitted on both loci at once, compared against the two specialist fits here, would show whether locus diversity buys robustness across the mismatch and what it costs on either diagonal (§6).

## 4 Scope and limitations

### What was evaluated

Two architectures were evaluated, but both are distributed by the same authors, share a training corpus, and have identical label spaces; the replication therefore controls for architecture and not for training data, label-space construction, or development group, and a model from an independent lineage might behave differently. Both were used as distributed, without fine-tuning on ITS2. A different classifier fitted on primer-bounded ITS2 does not show the collapse (§3.7), but that is evidence about what the deficiency is rather than a demonstration that fine-tuning these two models would remove it; our claim is about the pretrained artefacts practitioners would actually download rather than about the architectures in principle. Both were also run through their own distributed interface with default settings; we did not tune inference-time options and cannot exclude that some configuration would improve ITS2 performance, though the interface exposes no option relating to query length or amplicon region and issues no warning on short input, so the configuration we used is the one a practitioner following the documentation would arrive at.

### How the queries and strata were built

Novelty is defined by absence from the models’ label space, which reflects an earlier reference release, so a genus later split or renamed may be counted novel although the model encountered its sequences under a prior name; this biases the novel stratum towards better apparent classifier performance, making the observed degradation conservative. Of the 252 genera excluded as outside the models’ family space, about one in fifteen of those we could check carries an accepted name whose family the models do hold, so a small part of that exclusion reflects disagreement between the release’s taxonomy and an external nomenclator rather than genuine unplaceability (§2.2). One-per-genus sampling deliberately removes abundance weighting, so absolute values are not comparable with published abundance-weighted accuracies, only with each other; it also discards intrageneric variation, which for a multicopy region such as ITS is not negligible. Because it equalizes genera rather than families, the family-rank accuracies reported here remain weighted by how many genera each family contributes, and a macro-average over families would answer a different and equally reasonable question (§2.2); we report both weightings throughout (Appendix E). And disjoint record identifiers do not guarantee that no query has a nearly identical reference neighbour: three-fifths of the seen stratum has one at 99% identity or above, and although excluding them changes no conclusion (Appendix F), it does remove around four points of the reference-based methods’ seen-genus margin, so that margin is partly retrieval and should not be read as generalization alone (§2.4).

### What the scores do and do not mean

The MycoAI output used here as a confidence and novelty score is a softmax over the family label space. We used it ordinally, which is what AUROC and a threshold require, and additionally measured whether it is calibrated as a probability of correctness (Appendix G); we did not attempt to recalibrate either model, so the calibration reported is of the artefacts as distributed. The distinction between an input the models were not trained to receive in that format and a taxon they were never trained on runs through the whole ITS2 arm: on ITS2 both hold at once, so the novelty AUROC we report separates seen from novel genera within an unfamiliar input format rather than measuring whether the models detect that unfamiliarity (§3.5, Table 4). Novelty detection is in turn not the same as knowing whether a prediction is correct, and we measure only the first.

### What the ablation supports

The ablation establishing that the models weight the flanking regions above ITS2 uses constructed inputs, a padded fragment and an interphylum chimera, which no workflow would produce; they identify what the models’ predictions follow and are not estimates of performance on real data, and the padding arm additionally confounds absence of information with presence of misleading sequence. It localizes the dependence to the flanking regions as a block and no further: it does not separate ITS1 from 5.8S, does not identify which positions within them carry the effect, and says nothing about the internal mechanism, all of which would require position-resolved attribution (§6). Relatedly, following the flanks and classifying from the flanks are different properties and only the first is demonstrated. The account we give of why the two models differ in how strongly they follow the donor rests on their input representations rather than on an experiment: no arm manipulates sequence order while holding *k*-mer composition fixed, which is what would test it (§3.3).

### What was fitted once, or checked narrowly

The HiTaC results are single fits. Its estimator is deterministic given its training data (§2.8), so this bounds nothing about seeds, but it does leave the sensitivity of those fits to which reference records were sampled uncharacterized beyond the two depths tested. The robustness check on ITS2 delimitation compares two extractions agreeing on 99.7% of records, so it demonstrates independence of implementation rather than of methodology; a materially different ITS2 window might behave otherwise. The depth observation in §3.7 rests on two reference depths at one locus for a single classifier, enough to show that off-locus robustness is not fixed by architecture but not enough to characterise how it scales. And every arm of this study uses curated reference records, so sequencing error, primer bias and chimeras, all of which an environmental run produces, are absent throughout.

## 5 Reproducibility

Every value reported here is computed from per-query scored tables produced by a three-stage harness that constructs the query sets, references and ablation arms from UNITE, scores each method, and draws Figure 2 from the same summary table the text quotes, so figure and text cannot diverge. The per-rank label sets recovered from the distributed models are deposited as plain-text taxon lists, 3,695 genera and 791 families, so the stratification is reproducible without redistributing the models. Query sets, ablation arms and references are not deposited, being rebuilt from UNITE by those scripts; the pretrained classifiers and the UNITE release are cited below rather than redistributed. All analysis scripts, the per-query scored tables every value here is computed from, and the recovered label sets are available under the MIT license at https://github.com/ayobi/its-classifier-audit. The pinned software environment, the script inventory and representative commands are given in Appendix H.

## 6 Discussion

MycoAI is a substantial piece of work and the problem it addresses is real: assigning taxonomy to millions of ITS reads against a reference of comparable size is computationally awkward, and a model that classifies 10^5^ sequences in minutes is a genuine contribution to how such surveys are run. Nor is our finding that learned classification is the wrong approach: the best novel-genus recovery we measured came from a trained classifier, and it was trained in ten minutes on one percent of MycoAI’s data. Our finding is that the released MycoAI models are not ready for the data environmental mycology produces, and that the bar they fall below is lower than a comparison with alignment alone would suggest. On full-length reference sequences they are already outperformed by two long-standing reference-based methods at every rank and in both strata. On the ITS2 amplicon that environmental surveys overwhelmingly generate they lose roughly half their family accuracy in absolute terms, where those same methods lose under four points. Because records, stratification and reference rule were common to all five methods, the collapse cannot be attributed to the locus: ITS2 retains enough information for the other three methods to keep about 96% of their full-length seen-genus accuracy.

The ablation locates the failure precisely enough to say what would and would not address it. These models weight taxonomic information in the regions flanking ITS2 above information in ITS2 itself, to the point that supplying the flanks of an unrelated phylum causes the transformer to return that phylum’s family for most queries whose ITS2 is intact and correct. That is not a shortfall of context to be made up with more training data at the same locus. It is a dependence that predicts failure whenever the flanking regions are absent, which is the normal condition in amplicon sequencing. The larger model is the more dependent, following the donor roughly twice as strongly as the convolutional one, and the same direction appears within a single method in §3.7, where the deeper-trained HiTaC also lost more when moved off-locus. We read these together as suggesting that susceptibility grows with how much of the training record a model exploits, rather than as a claim about scaling, which two non-independent architectures cannot support. The constructive implication is that the deficiency is specific and addressable, and §3.7 demonstrates this rather than conjecturing it: the same hierarchical classifier, refitted on the ITS2 reference and given ITS2 queries, recovered seen-genus families at a rate that places it alongside best-hit alignment and SINTAX rather than beneath them, from a fit costing ten minutes on one percent of the MycoAI corpus. We did not fine-tune the MycoAI models themselves, so this establishes what the deficiency is rather than that it has been repaired in them, and our claim continues to concern the released artefacts rather than the architectures.

It is worth separating what these experiments demonstrate from what we take them to mean and from what remains conjecture, since the three are easy to run together. Demonstrated, on these queries: the two distributed models are outperformed by best-hit alignment, by SINTAX and by a ten-minute logistic regression at both loci and in both strata; the loss on ITS2 is an order of magnitude larger for them than for the other three; a classifier of the same fitted kind trained at the amplicon locus does not show that loss; and when the flanks of an unrelated phylum are supplied around an intact ITS2, the predictions follow the donor’s lineage rather than the query’s. Interpretation compatible with those observations, and the one we adopt: these models’ predictions are governed principally by information in the regions flanking ITS2, so removing those regions removes most of what they were using. Conjecture, which we flag as such: that ITS1 rather than 5.8S is the vehicle, which arm (iv) cannot separate; that the two models differ in how strongly they follow the donor because one retains sequence order and the other classifies an order-free 4-mer spectrum, which their documented input encodings make available but no arm here tests; that susceptibility grows with how much of the training record a model exploits, which two non-independent architectures and one auxiliary comparison cannot establish; and that models from other groups would behave the same way, which we did not test. Nothing here identifies the internal mechanism by which the flanking regions come to dominate, and following the flanks and classifying from the flanks remain different properties, of which only the first is shown.

The compounding problem is what the models report about themselves. On ITS2 neither score separates novel from known genera to any useful degree, one of the two sitting at chance, and a novelty flag calibrated on either detects genuine novelties at almost exactly the rate at which it fires on knowns. How that reaches a practitioner depends on the model, and the two cases are worth distinguishing because they fail differently (§3.5). The convolutional model is overconfident on this amplicon, so its output is confident names, mostly wrong, with nothing to indicate it. The transformer is not overconfident, and its score does track whether its own call is right; what it will not tell anyone is whether the genus was ever in the label space, and the price of using the score properly is discarding almost the whole run. Neither is a usable position, and neither is visible from a full-length evaluation. Three recommendations follow for how such models are reported. Accuracies should be given for the amplicon region the intended application generates, since here the two differ by a factor of four and the full-length figure is actively misleading about the ITS2 case. The length distribution of the training corpus should be stated, since it determines the range over which a reported accuracy can be expected to hold and is not currently recoverable from the model card. And a same-query classical baseline should be standard, because on this benchmark SINTAX and best-hit alignment were superior in every cell we measured, including on the novel genera that motivate learned approaches in the first place. That conclusion is not ours alone. In the benchmark table of a paper introducing a newer fungal foundation model, BLAST exceeds both MycoAI models at family and genus rank on the MycoAI test set [7], in the same work whose introduction attributes poor generalization to novel taxa to traditional methods. The baseline is often already present in these papers and simply not read as the comparison it constitutes.

For environmental mycology the practical consequence is narrow and immediate. A survey that amplifies ITS2 with primers in 5.8S and LSU delivers to these models exactly the input the ablation identifies as impoverished, and it does so for every sequence in the run rather than for an unlucky subset, so the failure is systematic rather than a tail. It is also silent: family calls are returned for everything, with confidence values that on this amplicon carry almost no information about which of them to trust. The resulting community table would be wrong in a structured way rather than a random one, since the errors are the models’ response to which flanks they were and were not given, and any ecology built on such a table would inherit that structure with nothing in the pipeline’s own output to flag it. The remedies available today are unglamorous and effective: classify ITS2 amplicons with a method fitted or referenced at that locus, and report a classical baseline computed on the same queries.

Two claims we might have made are not supported by our data, and we state them as such. Fixed label spaces are not the operative limitation: the reference-based methods consult a reference lacking the novel genus and are equally unable to name it, all five methods scoring exactly zero. And the classifiers’ weakness is not specific to novel taxa, since all five methods degraded by a similar 20 to 25 percentage points from seen to novel genera, making the learned representations uniformly weaker rather than differentially weak on dark matter.

What does distinguish the methods is whether their scores degrade informatively. Alignment identity and the SINTAX bootstrap remained strong novelty signals on ITS2; the classifiers’ scores fell to chance or within a whisker of it. A score that declines when the input is unfamiliar can be thresholded, calibrated, or used to abstain, and the machinery for turning such a score into a decision with a guaranteed error rate is the subject of a companion study [18]. A score that does not decline offers no purchase for any such mechanism, which is why we regard the loss of novelty signal on ITS2 as the more serious of the two failures reported here.

Several experiments would sharpen or challenge this account, and we list them in the order we would run them. Position-resolved attribution would replace a block-level ablation with a map: sliding-window occlusion across the full-length record, and for the transformer an attribution method such as integrated gradients, would show whether the dependence concentrates in ITS1, in 5.8S, or in neither, and attribution maps computed on the arm (iv) chimeras would compare the contribution of the query’s ITS2 against the donor’s flanks directly. Occlusion can return a flat profile even where a real dependence exists, in which case the negative result would bound the method rather than the hypothesis, so we would read it alongside the second experiment rather than instead of it. A cheaper arm would hold 4-mer composition fixed while permuting sequence order, which the convolutional model should by construction ignore and the transformer should not; that is the experiment our reading of their difference in §3.3 actually predicts, and it needs no attribution machinery. The second experiment measures the relation between truncation and representation directly: embedding full-length records and progressively truncated versions of the same records, and measuring how far the pair separates, would say how much of the ITS2 deficit is attributable to input format alone, which the present design confounds with absence of the flanks. Third, a classifier fitted jointly on full-length ITS and ITS2 and compared against the two specialist fits of §3.7 would show whether locus diversity buys robustness across the mismatch, and what it costs on either diagonal. Fourth, the comparison should eventually be run on real environmental reads rather than on reference records restricted to a subregion, since sequencing error, primer bias and chimeras are absent from every arm here. The same question can also be asked one level down, at the level of individual reads rather than of consensus sequences, where the same locus mismatch would be expected to appear.

### A Operational notes on the distributed packages

Two properties of the MycoAI package are met before a first prediction is possible and are recorded here for reproducibility, since a paper about what practitioners would download should say what downloading it entails. Neither affects any value reported in the main text, all of which was produced under the pinned environment of Appendix H. Both were encountered again, unchanged, when the checkpoints were reopened to confirm their configurations for §2.1, and we describe them as observed rather than as reported. First, mycoai/ init .py calls a telemetry service’s login routine at import, which raises when no credentials are present; because the checkpoints unpickle classes from that package, the call fires on torch.load as well as on direct import, and neither can proceed in a non-interactive shell until the routine is stubbed or the service placed in a disabled mode. Second, the checkpoints cannot be loaded by PyTorch 2.6 or later, in which torch.load defaults to refusing to unpickle arbitrary classes; under 2.13.0 this surfaces as an unpickling error naming SeqClassNetwork as a disallowed global, and loading requires passing weights_only=False or allowlisting that class, neither of which the package does.

A third message appears on every load and is benign, but is recorded here because the label-space recovery of §2.1 depends on the objects it concerns. The per-rank label encoders were serialized under scikit-learn 1.3.0 and are unpickled here under 1.3.2, which raises an inconsistent-version warning. The affected class stores its label inventory as a plain array, and the counts we recover from it reproduce MycoAI’s own published label file exactly at all six ranks, so the mismatch has no effect on the stratification.

One correction was likewise required for HiTaC. The reference was rewritten with a terminal semicolon, which its header parser expects and without which it silently truncates the final rank label by one character. Every family value reported in the main text is unchanged by that correction, family not being the terminal field.

### B Computation

All timings are on the same eight-thread CPU. Inference over the 5,222 queries took approximately 2 seconds for MycoAI-CNN and 295 seconds for MycoAI-BERT, a factor of about 140 between the two architectures. The two HiTaC fits, at 16,223 and 55,924 sequences, took approximately ten minutes each. No step in the study required a GPU, and the full set of arms reported here is reproducible in a few CPU-hours.

### C Robustness to the ITS2 delimitation: full values

Because the ITS2 arm inherits a boundary drawn by the release’s authors, ITS2 was independently re-extracted from the same queries and both versions classified on the identical 2,637 records for which re-extraction succeeded. The two delimitations proved close: 85.1% of records were identical and 99.7% identical or nested, with a median length difference of zero and medians of 167 and 165 bp. Both models were run on both extractions. Convolutional seen-genus family recovery was 18.7% and 18.3%, novel-genus 15.1% and 14.7%, coherence 51.5% and 51.4%; the transformer gave 28.6% and 28.4%, 18.4% and 18.5%, with coherence 56.3% and 56.5%. Neither model moves by more than 0.4 percentage points between delimitations, and the transformer moves less than the convolutional network, 0.2 and 0.1 points against 0.4 and 0.4. Flank dependence and boundary sensitivity are therefore not the same property: the model that follows the flanking regions more strongly (*§*3.3) is the less affected by where the ITS2 boundary is drawn, which is what one would expect if what matters is whether the flanks are present at all rather than their exact extent. The subset is also representative of the whole, recovery on it being close to the values obtained on all 5,222 queries for both models, 18.7% against 18.5% and 28.6% against 28.3%, so the records for which HMM anchoring succeeded are not appreciably easier. Because the two extractions are near-identical in sequence, this establishes independence of implementation rather than of methodology, and does not exclude the possibility that a substantially different ITS2 window would behave differently.

### D Sensitivity to reference depth

Because the reference available to the reference-based methods is a design choice, the full-ITS alignment arm was repeated at three per-genus depths (Table 7). Seen-genus recovery rose monotonically with depth, from 90.1% to 93.8%, as did genus recovery on that stratum (77.9% to 87.3%) and novelty separability (AUROC 0.836 to 0.887). Novel-genus recovery did not: it peaked at twenty sequences per genus (67.2%) and fell at fifty (64.3%).

**Table 7:** Sensitivity of the full-ITS alignment arm to reference depth. The classifiers do not consult the reference and their results are unchanged throughout.

| cap per genus | reference size | family, seen | family, novel | AUROC |
| --- | --- | --- | --- | --- |
| 5 | 16,352 | 90.1% | 66.1% | 0.836 |
| 20 | 56,327 | 92.3% | 67.2% | 0.872 |
| 50 | 113,455 | 93.8% | 64.3% | 0.887 |

Alignment exceeds both classifiers on both strata at every depth tested.

The asymmetry follows from what depth can supply. A seen-genus query gains conspecifics and congeners as the reference deepens, so its best hit improves; a novel-genus query cannot gain a same-genus relative at any depth, while the growing reference offers more opportunities for a spurious best hit from the wrong family. Deeper references therefore assist known taxa and mildly impede novel ones. For the present purpose the relevant observation is that alignment exceeded both classifiers on both strata at every depth tested, so the ranking does not depend on this choice, and that the twenty-per-genus reference used throughout is close to optimal for the novel-genus stratum rather than generous to alignment.

### E Family recovery macro-averaged over families

Because queries are drawn one per genus (§2.2), a family contributes as many queries as it has genera, and the pooled family-rank accuracies in Table 1 are weighted accordingly. The seen stratum spans 732 families and the novel stratum 432, with a median of two genera per family and a maximum of 68 and 77; pooling therefore gives a few large families considerable influence. Table 8 repeats every family-rank figure macro-averaged over families, each family counting once regardless of how many genera represent it.

**Table 8:** Family recovery pooled over queries, as in Table 1, and macro-averaged over families. Macro-averaging lowers every method, since small families are placed less accurately than large ones, but it does not alter the separation the paper rests on.

| locus | method | seen genus |  | novel genus |  |
| --- | --- | --- | --- | --- | --- |
|  |  | pooled | macro | pooled | macro |
| full ITS | alignment | 92.3% | 89.3% | 67.2% | 53.9% |
|  | SINTAX | 92.0% | 88.0% | 70.3% | 55.3% |
|  | HiTaC | 92.3% | 88.5% | 71.4% | 56.5% |
|  | MycoAI-BERT | 77.9% | 68.7% | 57.7% | 42.9% |
|  | MycoAI-CNN | 76.5% | 66.3% | 54.3% | 38.5% |
| ITS2 | alignment | 88.6% | 86.7% | 60.5% | 49.1% |
|  | SINTAX | 88.5% | 84.9% | 63.6% | 49.4% |
|  | HiTaC | 89.2% | 84.9% | 62.5% | 45.9% |
|  | MycoAI-BERT | 28.3% | 16.4% | 18.8% | 11.2% |
|  | MycoAI-CNN | 18.5% | 14.7% | 13.8% | 9.0% |

At full length the two pretrained models lose more under macro-averaging than the other three on seen genera, 9.2 and 10.1 points against 3.0 to 4.0, so their pooled figures are the more generous of the two and the gap widens when families are weighted equally. On novel genera every method loses 11 to 17 points and the ordering is preserved. One ordering does change. The three leading methods are separated on ITS2 seen genera by 0.7 points pooled, with HiTaC highest; macro-averaged they are separated by 1.8 points with alignment highest and HiTaC level with SINTAX. Since McNemar’s test does not distinguish these three at either locus (§3.1), we take the reversal as further reason not to interpret gaps of that size rather than as a result about HiTaC.

### F The comparison with near-duplicate queries removed

Queries and reference share no record identifiers (*§*2.4), but a query may still have a reference neighbour that is nearly or exactly identical, and a method credited for placing such a query is being credited for retrieval rather than generalization. Best-hit percentage identity from the alignment arm measures this directly. The two strata are far apart on it: at full length the median seen-genus query sits at 99.4% identity to its nearest reference neighbour and the median novel-genus query at 85.5%, with 59.3% of the seen stratum at 99% or above and 26.5% matching a reference exactly. The ITS2 figures are similar, and the proportion of exact matches is higher (48.3%), as expected when a shorter window is compared.

Table 9 recomputes the comparison after discarding, at each locus, every query whose best reference hit reaches 99% identity. This removes three-fifths of the seen stratum and under a tenth of the novel one.

**Table 9:** Family recovery on all queries and on the subset whose nearest reference neighbour lies below 99% identity. The same query subset is used for every method at a locus, so the columns remain comparable.

| locus | method | seen genus |  | novel genus |  |
| --- | --- | --- | --- | --- | --- |
|  |  | all | < 99% | all | < 99% |
| full ITS | alignment | 92.3% | 87.7% | 67.2% | 66.9% |
|  | SINTAX | 92.0% | 88.2% | 70.3% | 70.3% |
|  | HiTaC | 92.3% | 88.5% | 71.4% | 71.1% |
|  | MycoAI-BERT | 77.9% | 77.2% | 57.7% | 56.3% |
|  | MycoAI-CNN | 76.5% | 75.6% | 54.3% | 52.7% |
| ITS2 | alignment | 88.6% | 84.2% | 60.5% | 59.7% |
|  | SINTAX | 88.5% | 84.4% | 63.6% | 63.0% |
|  | HiTaC | 89.2% | 82.8% | 62.5% | 61.4% |
|  | MycoAI-BERT | 28.3% | 26.0% | 18.8% | 17.0% |
|  | MycoAI-CNN | 18.5% | 17.9% | 13.8% | 13.3% |

Subset sizes: 1,355 seen and 1,731 novel queries at full length, 1,253 and 1,714 on ITS2.

**Table 10:** Calibration of each score as a probability that the family returned is correct, pooled across strata. ECE and MCE are the mean and maximum absolute gap between reported score and observed accuracy across fifteen bins; “mean score *−* accuracy” is the aggregate over- or under-confidence.

| locus | method | mean | accuracy | mean score | ECE | MCE |
| --- | --- | --- | --- | --- | --- | --- |
|  |  | score |  | – accuracy |  |  |
| full ITS | SINTAX | 0.852 | 0.842 | +0.011 | 0.055 | 0.166 |
|  | MycoAI-BERT | 0.769 | 0.706 | +0.063 | 0.070 | 0.097 |
|  | MycoAI-CNN | 0.678 | 0.684 | –0.007 | 0.065 | 0.119 |
| ITS2 | SINTAX | 0.863 | 0.795 | +0.068 | 0.071 | 0.125 |
|  | MycoAI-BERT | 0.252 | 0.249 | +0.003 | **0.019** | 0.177 |
|  | MycoAI-CNN | 0.394 | 0.168 | +0.225 | **0.226** | 0.546 |

Best-hit identity is omitted: it is a sequence similarity rather than a probability and is not offered as one, and treating it as such gives an expected calibration error of 0.090 at full length and 0.117 on ITS2, which measures the units rather than the method.
Brier scores, in the same row order: 0.093, 0.152, 0.159 at full length and 0.114, 0.151, 0.193 on ITS2. The Brier score conflates calibration with discrimination and is reported for completeness.

### G Calibration of the scores used as confidence

The manuscript uses each method’s score ordinally, which is what AUROC and a threshold require (*§*2.5). The distributed interface nonetheless returns that score to users alongside a name, which invites reading it as the probability that the name is right. This appendix asks whether it can be read that way. For each method we compare the score with the observed accuracy of the top-ranked family call in fifteen equal-width bins, and summarize the gap as expected calibration error, its worst bin as maximum calibration error, and the mean squared difference between score and outcome as a Brier score.

Two results follow, and they concern different models. MycoAI-CNN is substantially over-confident on ITS2 and only there: its expected calibration error rises from 0.065 at full length to 0.226, its worst bin is off by 0.546, and thresholding does not rescue it, since the queries it scores above 0.9 are correct 49.8% of the time and those above 0.8 only 43.7%. MycoAI-BERT is not overconfident on ITS2 at all; its expected calibration error of 0.019 is the lowest value in the table, and the reason is that its ITS2 scores are simply low, with a median of 0.171 against 0.322 for the convolutional model. Thresholding it at 0.8 retains 3.3% of queries at 78.7% accuracy, against 65.1% of queries at 86.0% accuracy for the same model and threshold at full length.

Calibration and discrimination are therefore separable properties, and MycoAI-BERT on ITS2 separates them about as far as they go: a score that is well calibrated in aggregate (Table 10) and close to uninformative about whether the query’s genus is novel (AUROC 0.541, *§*3.5). A score reporting a quarter and correct a quarter of the time is calibrated by construction, and a practitioner cannot act on it. Nothing in this appendix is a recalibration; both models are assessed as distributed.

### H Software environment, harness and commands

Analyses used vsearch v2.31.0, ITSx v1.1.3, ITSxRust v0.2.2, and mycoai-its v0.0.5 with torch v2.13.0, scikit-learn v1.3.2, numpy v1.26.4 and biopython v1.87 on Python 3.10.20.

The exclusion costs the three leading methods 3.8 to 6.3 points of seen-genus recovery and the two pretrained models 0.6 to 2.3, which is the expected direction: a method that consults a reference gains most from a near-identical neighbour and a fixed-label classifier gains least. The leaders’ full-length margin over the pretrained models narrows from about fifteen points to about eleven and does not close. On the novel stratum, where near-duplicates are rare by construction, nothing moves by more than 1.6 points. Repeating the exclusion at 97% identity, which removes about three-quarters of the seen stratum, moves the leaders a further two to four points and leaves the same ordering.

Several of these are pinned by MycoAI’s dependency constraints rather than being current releases, scikit-learn in particular being held at the 1.3 series, so they are reported as installed rather than as recommended. Because the torch release used postdates the change described in Appendix A, every classification reported here was produced with the unpickling default overridden; that override is the only modification made to the distributed inference path, and it changes what can be loaded rather than what is computed. The harness is organised in three stages:

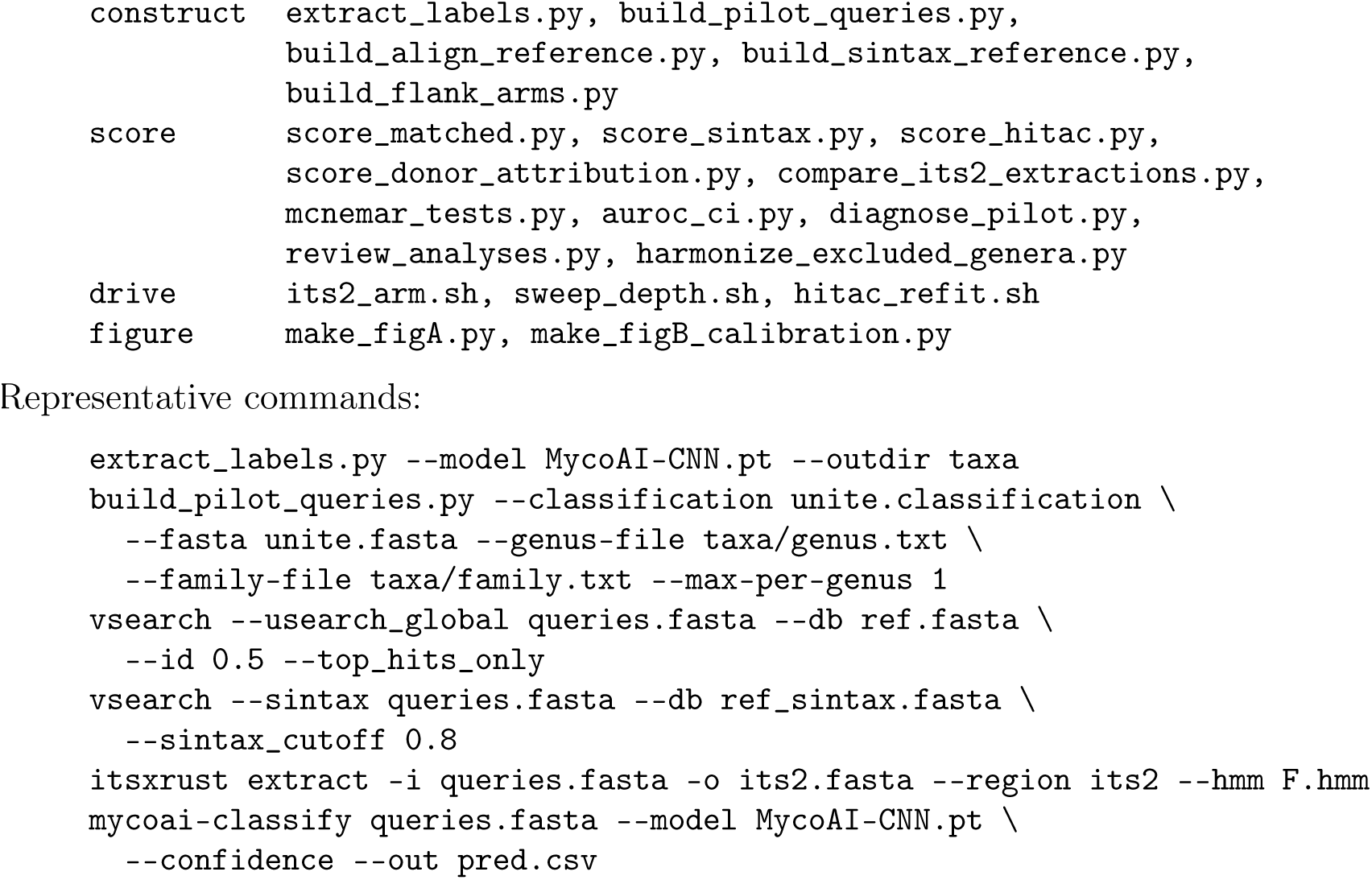

## Author contributions

Aaron O’Brien conceived the study, built the harness, ran all analyses and wrote the original draft. Auguste Gardette critically reviewed the manuscript and contributed to the framing of the evaluation and to the interpretation of the results, in particular the treatment of method terminology, the distinction between input-format and taxonomic novelty, and the attribution and calibration analyses set out in the Discussion. César Marín critically reviewed and revised the manuscript, contributing edits throughout and additions to the literature cited. Pilar Parada acquired the funding (CORFO 23PTECCC-247149) and supervised the project. All authors read and approved the submitted version.

## Competing interests

Two of the authors of this manuscript, Aaron O’Brien and Pilar Parada, are authors of ITSxRust [19], which Section 3.4 uses solely as an independent re-implementation for a robustness check. The reported result there is that the two extractions agree to within 0.4 percentage points for either model, which is neutral with respect to both tools, and the conclusion of that section would be unchanged had the re-extraction been performed with any other implementation.

## Data and code availability

Harness scripts, the recovered label-space taxon lists, the per-query scored tables and the results summary are available at https://github.com/ayobi/its-classifier-audit under the MIT license. Reference data: UNITE+INSD 2024 via Zenodo 13336328. Pretrained classifiers: Zenodo record 10904344.

## Acknowledgements

This work was supported by CORFO grant 23PTECCC-247149 to Pilar Parada. César Marín thanks Fondecyt Regular Project No. 1240186 (ANID, Chile). We thank the Centro de Biotecnología de Sistemas for compute resources.

## Use of generative AI

Anthropic’s Claude and GitHub Copilot were used in preparing this work: to draft and revise manuscript text, to write and debug the analysis and figure scripts deposited with it, and to check bibliographic details. All analyses were run by the authors, all reported values were verified against the primary output, and the authors are responsible for the content of the manuscript.

## Notes

https://github.com/ayobi/its-classifier-audit

### Competing Interest Statement

Both authors are authors of ITSxRust (bioRxiv 2026.02.25.707950), which is used in this manuscript solely as an independent re-implementation for a robustness check. The reported result is that the two extractions agree, which is neutral with respect to both tools. The authors declare no financial competing interests.

### Summary of Updates

Auguste Gardette critically reviewed the manuscript and contributed to the framing of the evaluation and to the interpretation of the results, in particular the treatment of method terminology, the distinction between input-format and taxonomic novelty, and the attribution and calibration analyses set out in the Discussion.

